# Antagonistic control of DDK binding to licensed replication origins by Mcm2 and Rad53

**DOI:** 10.1101/2020.05.04.077628

**Authors:** Syafiq Abd Wahab, Dirk Remus

**Author notes:** Corresponding author: Molecular Biology Program, Memorial Sloan Kettering Cancer Center, 1275 York Avenue, New York, NY 10065, USA.

## Abstract

Eukaryotic replication origins are licensed by the loading of the replicative DNA helicase, Mcm2-7, in inactive double hexameric form around DNA. Subsequent origin activation is under control of multiple protein kinases that either promote or inhibit origin activation, which is important for genome maintenance. Using the reconstituted budding yeast DNA replication system, we find that the flexible N-terminal tail of Mcm2 promotes the stable recruitment of Dbf4-dependent kinase (DDK) to Mcm2-7 double hexamers, which in turn promotes DDK phosphorylation of Mcm4 and -6 and subsequent origin activation. Conversely, we demonstrate that the checkpoint kinase, Rad53, inhibits DDK binding to Mcm2-7 double hexamers. Unexpectedly, this function is not dependent on Rad53 kinase activity, but requires Rad53 activation by trans-autophosphorylation, suggesting steric inhibition of DDK by activated Rad53. These findings identify critical determinants of the origin activation reaction and uncover a novel mechanism for checkpoint-dependent origin inhibition.

## Introduction

To ensure the timely, accurate, and complete duplication of their genomes prior to cell division, eukaryotic cells initiate DNA replication at many replication origins distributed along the length of each chromosome. From each origin two replication forks emanate in opposite direction to form a replication bubble. Although bidirectional origin firing has long been recognized as a universal feature of chromosomal DNA replication in all domains of life (Huberman and Riggs, 1968; Prescott and Kuempel, 1972), how pairs of oppositely oriented replication forks are established at chromosomal origins remains poorly understood. In eukaryotes, two copies of the replicative DNA helicase, Mcm2-7, are loaded as a stable double-hexameric complex around double-stranded DNA (dsDNA) at the origin (Evrin et al., 2009; Miller et al., 2019; Remus et al., 2009). Mcm2-7 comprise six related proteins of the AAA+ family of ATPases that assemble into a hexameric ring with defined subunit order. Intriguingly, the two hexamers in a Mcm2-7 double hexamer (DH) associate in a head-to-head configuration, thus providing a platform for the establishment of oppositely oriented sister replisomes. (Abid Ali et al., 2017; Evrin et al., 2009; Li et al., 2015; Miller et al., 2019; Noguchi et al., 2017; Remus et al., 2009). However, Mcm2-7 DHs are catalytically inactive and require the regulated association of the essential helicase co-factors Cdc45 and GINS to form two active replicative DNA helicase complexes, termed CMG (<u>C</u>dc45-<u>M</u>CM-<u>G</u>INS), which encircle single-stranded DNA (ssDNA) during unwinding (Bell and Labib, 2016). Therefore, to ensure bidirectional origin firing, mechanisms must exist that ensure simultaneous progression of both helicase complexes from the origin. As the head-to-head orientation of CMG helicases assembled at the origin requires them to pass each other during origin activation, it has been proposed that an active CMG encircling ssDNA is blocked by a dsDNA-encircling inactive CMG potentially formed around the opposite Mcm2-7 hexamer, thereby imposing origin bidirectionality (Douglas et al., 2018; Georgescu et al., 2017).

The replication of chromosomes from multiple replication origins necessitates strict control mechanisms that prevent origins from re-firing within one cell cycle in order to maintain genome stability. Such re-replication control is achieved by a two-step mechanism that temporally separates helicase loading in late M and G1 phase from helicase activation in S phase (Bell and Labib, 2016). Helicase activation is controlled by two cell cycle-regulated protein kinases, Dbf4-dependent kinase (DDK) and cyclin-dependent kinase (CDK), which act in conjunction with a defined set of co-factors, comprising Sld3·7, Sld2, Dpb11, Pol ε, and Mcm10, in addition to GINS and Cdc45, to mediate CMG assembly (Douglas et al., 2018). The essential targets of CDK during CMG assembly are Sld2 and Sld3, which physically interact with distinct Dpb11 BRCT domains when phosphorylated to recruit GINS and Pol ε to the origin (Bell and Labib, 2016). The mechanism by which DDK promotes CMG assembly is somewhat obscure. Genetic and biochemical studies demonstrate that Mcm2-7 are the essential targets for DDK during DNA replication (Deegan et al., 2016; Hardy et al., 1997; Randell et al., 2010; Sheu and Stillman, 2010; Yeeles et al., 2015). Accordingly, DDK phosphorylation of the flexible N-terminal tails of Mcm4 and -6 promotes the recruitment of Sld3·7 to Mcm2-7 DHs by generating phosphorylation-dependent binding sites for Sld3, which in turn recruits Cdc45 (Deegan et al., 2016; Gros et al., 2014; Heller et al., 2011; Tanaka et al., 2011; Yeeles et al., 2015). However, the nature and stoichiometry of the Sld3 interaction with Mcm2-7 DHs is unclear, as Sld3 exhibits limited sequence specificity for phospho-dependent binding sites, and phospho-mimicking mutations in either Mcm4 or -6 are sufficient to bypass the requirement for DDK (Deegan et al., 2016; Randell et al., 2010). Moreover, other mutations in Mcm5 and Mcm4 that do not involve phospho-mimetic amino acid substitutions can also bypass the requirement for DDK (Hardy et al., 1997; Sheu and Stillman, 2010). It is not known, if these mutations allow Sld3 recruitment in the absence of Mcm2-7 phosphorylation, or if they bypass the requirement for Sld3 altogether.

Replication origins do not fire simultaneously upon S phase entry, but in a staggered programmatic manner throughout S phase (Rhind and Gilbert, 2013). The replication timing program is in part imposed by limiting concentrations of the initiation factors Sld2, Dpb11, Sld3, Cdc45, and Dbf4, which restrict CMG assembly to subsets of origins throughout S phase (Mantiero et al., 2011; Tanaka et al., 2011). While limiting initiation factors that interact transiently with origins are thought to be recycled from early to late origins in a normal S phase, such recycling is prevented by the S phase checkpoint to inhibit late origin firing. The S phase checkpoint is a vital kinase signaling cascade that induces a range of cellular responses in addition to the inhibition of origin firing, including increasing dNTP levels, stabilization of stalled replication forks, and DNA repair to promote genome maintenance during replication stress (Pardo et al., 2017).

In budding yeast, the minimal targets for checkpoint-dependent origin inhibition are Sld3 and Dbf4, which are substrates for the checkpoint effector kinase, Rad53 (Lopez-Mosqueda et al., 2010; Zegerman and Diffley, 2010). Rad53 phosphorylation of Sld3 inhibits its physical interactions with Mcm2-7, Cdc45, and Dpb11 (Deegan et al., 2016; Lopez-Mosqueda et al., 2010; Zegerman and Diffley, 2010). How Rad53 inhibits Dbf4 is not clear. Rad53 was found to inhibit DDK kinase activity *in vitro*, but the mechanism of inhibition is unknown (Kihara et al., 2000; Weinreich and Stillman, 1999). Moreover, DDK activity in these studies was assessed using isolated Mcm2 or Mcm7 subunits as substrates or by measuring DDK autophosphorylation (Kihara et al., 2000; Weinreich and Stillman, 1999). However, since DDK exhibits high specificity for Mcm4 and -6 in the context of Mcm2-7 DHs, the effect of Rad53 on DDK activity at licensed origins remains to be determined (Francis et al., 2009; Randell et al., 2010; Sheu and Stillman, 2006; Sun et al., 2014). Rad53 has also been reported to disrupt DDK-chromatin association in cells treated with hydroxyurea (HU) (Pasero et al., 1999). DDK binds to chromatin via an interaction with Mcm2-7 at licensed replication origins (Dowell et al., 1994; Francis et al., 2009; Jares and Blow, 2000; Jares et al., 2004; Sato et al., 2003; Sheu and Stillman, 2006; Takahashi and Walter, 2005; Weinreich and Stillman, 1999; Yanow et al., 2003). Several potential DDK docking sites have been identified in Mcm2-7 based on pairwise interaction studies with individual Mcm2-7 subunits (Ramer et al., 2013; Sheu and Stillman, 2006). However, it has not been tested to what extent these contribute to DDK binding in the context of the Mcm2-7 DH, which is known to greatly stimulate DDK substrate specificity and kinase efficacy (Francis et al., 2009; Sun et al., 2014). How Rad53 may affect the DDK- chromatin association has not been addressed.

Here we employ the reconstituted budding yeast DNA replication system to investigate the mechanism of DDK docking to Mcm2-7 DHs and its regulation by Rad53 (Devbhandari et al., 2017; Devbhandari and Remus, 2020). We find that the flexible Mcm2 N-terminal tail is necessary for DDK docking onto Mcm2-7 DHs, Mcm4 and -6 phosphorylation, and origin firing. The data suggests that the N-terminal tails of both Mcm2 protomers in the Mcm2-7 DH are required for efficient origin activation. Intriguingly, Rad53 disrupts DDK binding to Mcm2-7 DHs in a manner that is independent of Rad53 kinase activity, but dependent on prior activation of Rad53. These observations have important implications for the mechanism of bidirectional origin firing and its regulation by the replication checkpoint.

## Results

### Residues 1-127 of Mcm2 are dispensable for Mcm2-7 DH stability

While the structured N-terminal domains (NTDs) and AAA+ ATPase domains are conserved between archaeal and eukaryotic MCM proteins, the long unstructured N-terminal tails of Mcm2, -4, and -6 are unique to eukaryotes. The N-terminal tails of Mcm4 and -6 play a fundamental role during DNA replication by serving as phospho-acceptors for DDK kinase during origin activation (Deegan et al., 2016; Francis et al., 2009; Randell et al., 2010; Sheu and Stillman, 2006). In contrast, the Mcm2 N-terminal tail contains a nuclear localization sequence (NLS) that mediates nuclear import of Cdt1×Mcm2-7(Liku et al., 2005) and a conserved histone H3/H4 binding domain (HBD) that controls nucleosome segregation at replication forks (Foltman et al., 2013; Gan et al., 2018; Huang et al., 2015; Petryk et al., 2018). As nuclear import is irrelevant for DNA replication with purified proteins, and chromatin can be omitted from DNA replication reactions *in vitro*, we asked whether the Mcm2 N-terminal tail possesses a basic DNA replication function. For this we engineered a TEV protease cleavage site downstream of the HBD, between residues A127 and Y128 (Mcm2-TEV; **Figure 1A**). Purified Cdt1×Mcm2-7 complexes harboring Mcm2-TEV (Cdt1×MCM^2-TEV^) are quantitatively and specifically cleaved at the engineered TEV site by TEV protease, resulting in Cdt1×Mcm2-7 complexes containing the Mcm2-Δ127 N-terminal truncation, whereas wildtype Cdt1×Mcm2-7 complexes are resistant to TEV protease cleavage (**Figure 1B, Supplementary Figure 1**).

**Figure 1.**
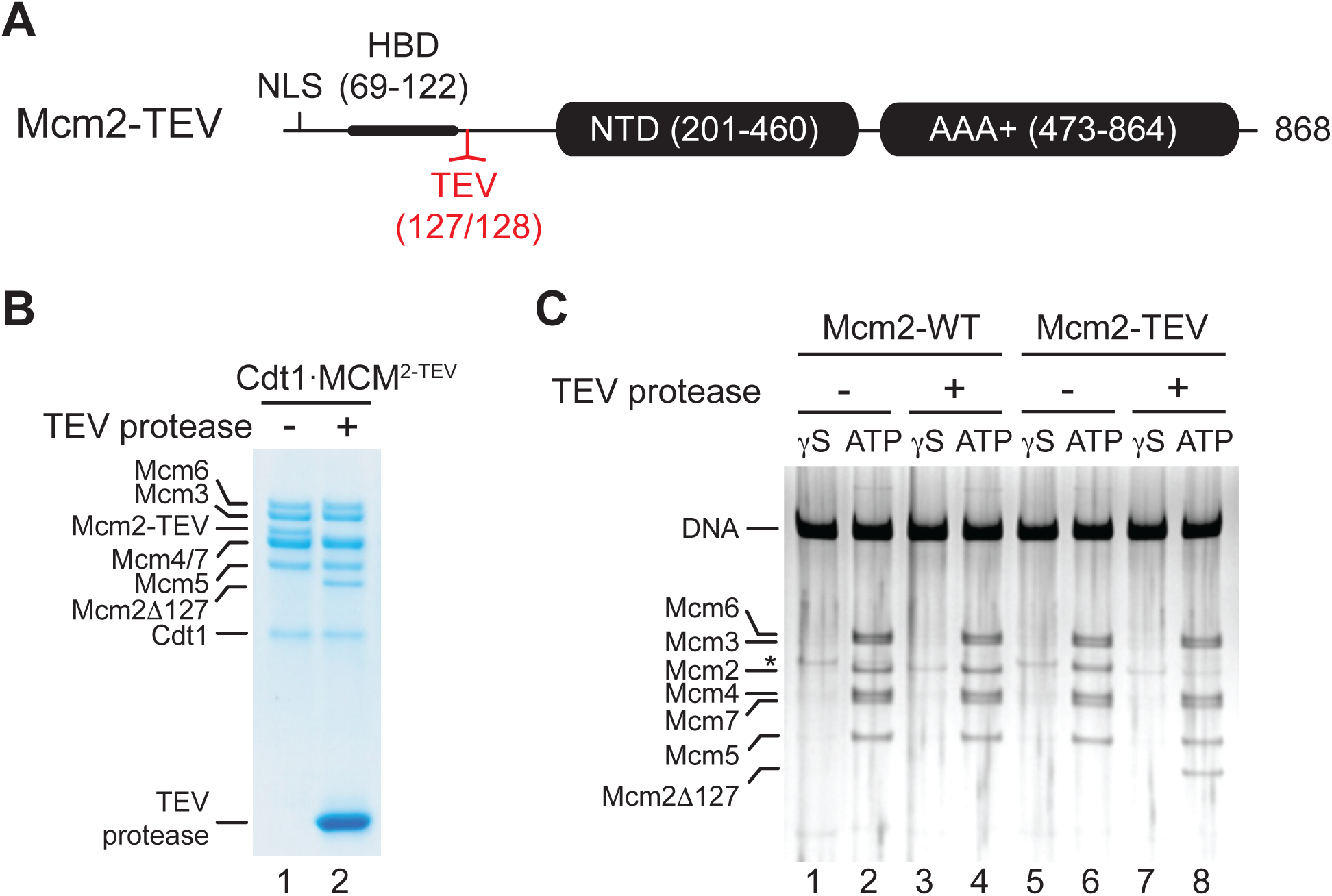
Residues 1-127 of the Mcm2 N-terminal tail are dispensable for Mcm2-7 DH stability. (**A**) Schematic of Mcm2 domain structure. Numbers indicate amino acid positions. The position of the TEV cleavage site is highlighted in red. NLS: Nuclear localization sequence; HBD: Histone binding domain; NTD: N-terminal domain; AAA+: ATPase domain. (**B**) Cdt1·MCM^2-TEV^ was mock-treated or digested with TEV protease for 1 hour at 30°C, as indicated. Reactions were fractionated on SDS-PAGE and stained with Coomassie blue. (**C**) Mcm2-7 loading reactions were performed on 3 kbp ARS305-containing DNA in the presence of ATPγS (γS) or ATP as indicated. DNA-bound material was washed with high-salt buffer, mock-treated or digested with TEV protease as indicated, washed again with high-salt buffer, and analyzed by SDS-PAGE and silver staining.

First, we tested if proteolytic truncation of the Mcm2 N-terminal tail affects Mcm2-7 DH stability. For this we performed reconstituted Mcm2-7 loading reactions on origin-containing DNA using purified ORC, Cdc6, and either wildtype Cdt1×MCM or Cdt1×MCM^2-TEV^ (Remus et al., 2009). Following Mcm2-7 loading, DNA-bound complexes were washed to remove free proteins, treated with TEV protease, and analyzed by SDS-PAGE. The DNA was immobilized on paramagnetic streptavidin-coated beads via a single photo-cleavable 5’-terminal biotin moiety to allow elution of the DNA from the beads with UV light for analysis. To differentiate Mcm2-7 DHs loaded around DNA from potential other more loosely associated complexes, such as the OCCM (<u>O</u>RC- <u>C</u>dc6-<u>C</u>dt1-<u>M</u>CM), DNA-bound complexes were washed with a high-salt buffer prior to DNA elution (Remus et al., 2009; Yuan et al., 2017). While Mcm2-7 DHs are resistant to salt-elution from the DNA, loading factors and loosely associated Mcm2-7 complexes are efficiently disrupted by stringent salt washes. In addition, control Mcm2-7 loading reactions were carried out in the presence of non-hydrolyzable ATPγS as Mcm2-7 DH formation is strictly dependent on ATP hydrolysis (Bell and Labib, 2016; Remus et al., 2009). Using this approach, we demonstrate comparable DNA loading efficiencies for wildtype Cdt1×MCM and Cdt1×MCM^2-TEV^. Moreover, truncation of the Mcm2 N-terminal tail from Mcm2-TEV-containing DHs, despite being efficient, did not negatively affect Mcm2-7 DH retention on DNA (**Figure 1C**). Thus, maintenance of Mcm2-7 DHs is not dependent on residues 1-127 of the Mcm2 N-terminal tail.

### Residues 1-127 of Mcm2 are important for DNA replication

Next we tested if residues 1-127 of the Mcm2 N-terminal tail are required for DNA replication *in vitro*. For this we performed DNA replication reactions both on naked DNA templates and reconstituted chromatin as described previously (Devbhandari et al., 2017). As FACT and Nhp6 have been demonstrated to promote replisome progression through chromatin *in vitro*, purified FACT and Nhp6 were also included in chromatin replication reactions here (**Supplementary Figure 2**) (Kurat et al., 2017). TEV protease cleavage of the Mcm2 N-terminal tail was induced following Mcm2-7 loading (**Figure 2A**).

**Figure 2.**
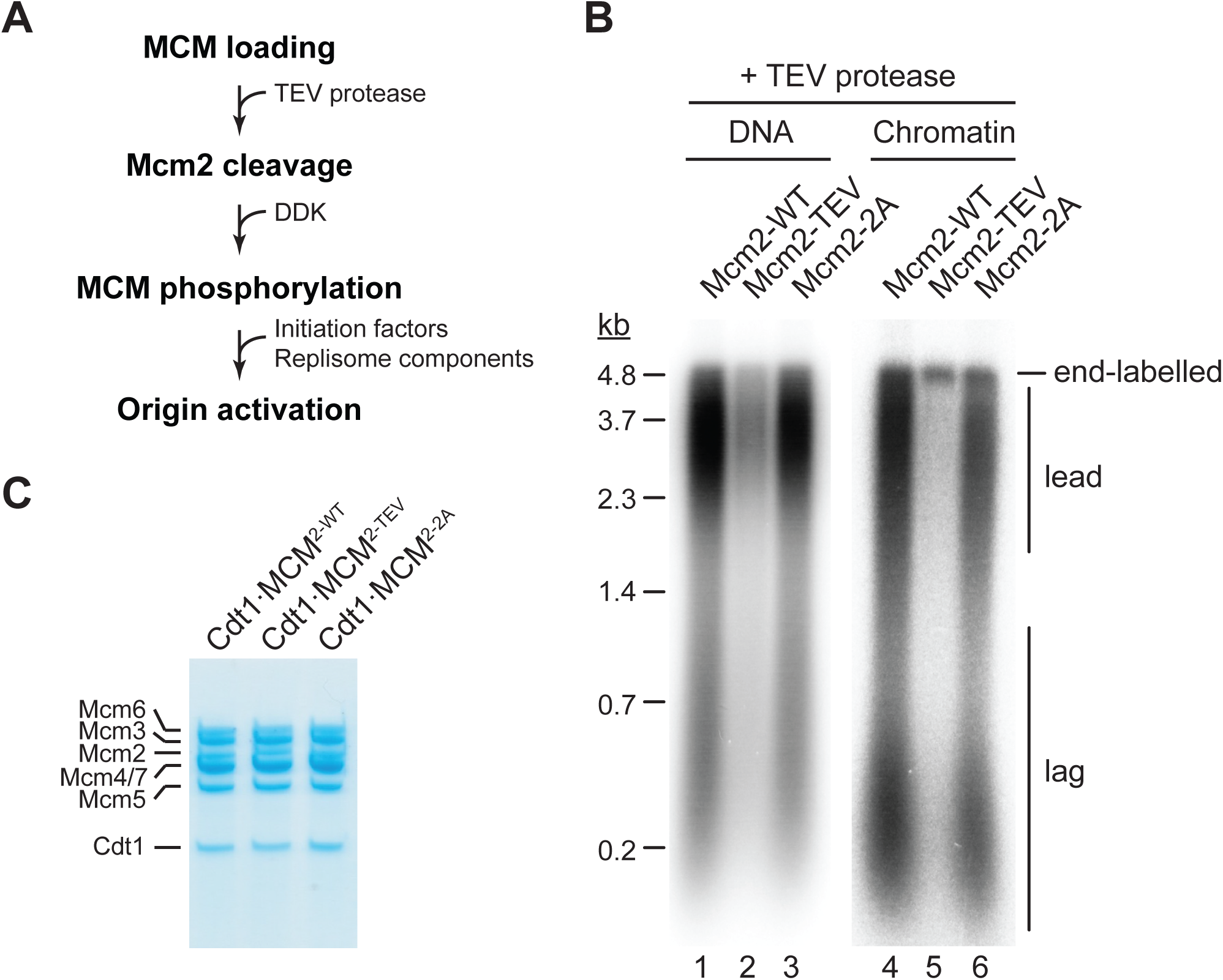
The Mcm2 N-terminal tail is important for DNA replication. (**A**) Experimental outline. (**B**) *In vitro* DNA replication reactions were performed on naked (lanes 1-3) or chromatinized (lanes 4-6) circular plasmid DNA (p1017, 4.8 kbp). TEV protease was added to each reaction following Mcm2-7 loading for 1 hour at 30°C, before addition of DDK and standard initiation / replisome factors. Chromatin replication reactions additionally contained FACT and Nhp6. Products were analyzed by 0.8 % denaturing agarose gel-electrophoresis and autoradiography. Lead: Leading strand product; lag: Lagging strand product. (**C**) Purified Cdt1·MCM complexes containing either wildtype Mcm2 (Cdt1·MCM^2-WT^), Mcm2-TEV (Cdt1·MCM^2-TEV^), or Mcm2-2A (Cdt1·MCM^2-2A^).

Intriguingly, truncation of Mcm2 residues 1-127 largely attenuated DNA replication (**Figure 2B**). Inhibition of DNA replication in the presence of Mcm2-TEV was dependent on TEV protease (**Supplementary Figure 3**). The DNA replication defect was not due to a loss of HBD function, as mutation of two conserved tyrosine residues, Y82 and Y91, in the Mcm2 HBD (Cdt1×MCM^2-2A^, **Figure 2C**) that have been previously shown to disrupt histone H3/H4 and FACT binding to Mcm2 has little effect on DNA replication using either DNA or chromatin as a template (Foltman et al., 2013). This is consistent with previous reports demonstrating that yeast cells harboring the *mcm2-2A* allele are viable and exhibit only mild replication defects (Foltman et al., 2013). This data reveals that the Mcm2 N-terminal tail performs a fundamental function during normal DNA replication that is distinct from its histone H3/H4 chaperone activity.

### The Mcm2 N-terminal tail promotes DDK function during the initiation of DNA replication

In order to dissect which step in the DNA replication reaction is defective in the absence of residues 1- 127 of Mcm2 we performed order-of-addition experiments by adding TEV protease at various steps of the origin firing pathway (**Figure 3A**). In control experiments, TEV protease did not disrupt the DNA replication proficiency of wild-type Cdt1×MCM when added prior to DDK immediately after Mcm2-7 loading (**Figure 3B**). Conversely, purified Cdt1×MCM^2Δ127^, obtained by prior TEV protease digestion of Cdt1×MCM^2-TEV^, was deficient for DNA replication, as expected (we show below that the replication defect is not due to a MCM loading defect). As before, in the presence of Cdt1×MCM^2-TEV^, addition of TEV protease to the reaction immediately after Mcm2-7 loading inhibited origin activation. In striking contrast, addition of TEV protease after DDK or Sld3×7 allowed DNA replication to proceed normally. This data indicates that Mcm2 residues 1-127 promote DDK function during the initiation of DNA replication but are dispensable at later steps of the DNA replication reaction.

**Figure 3.**
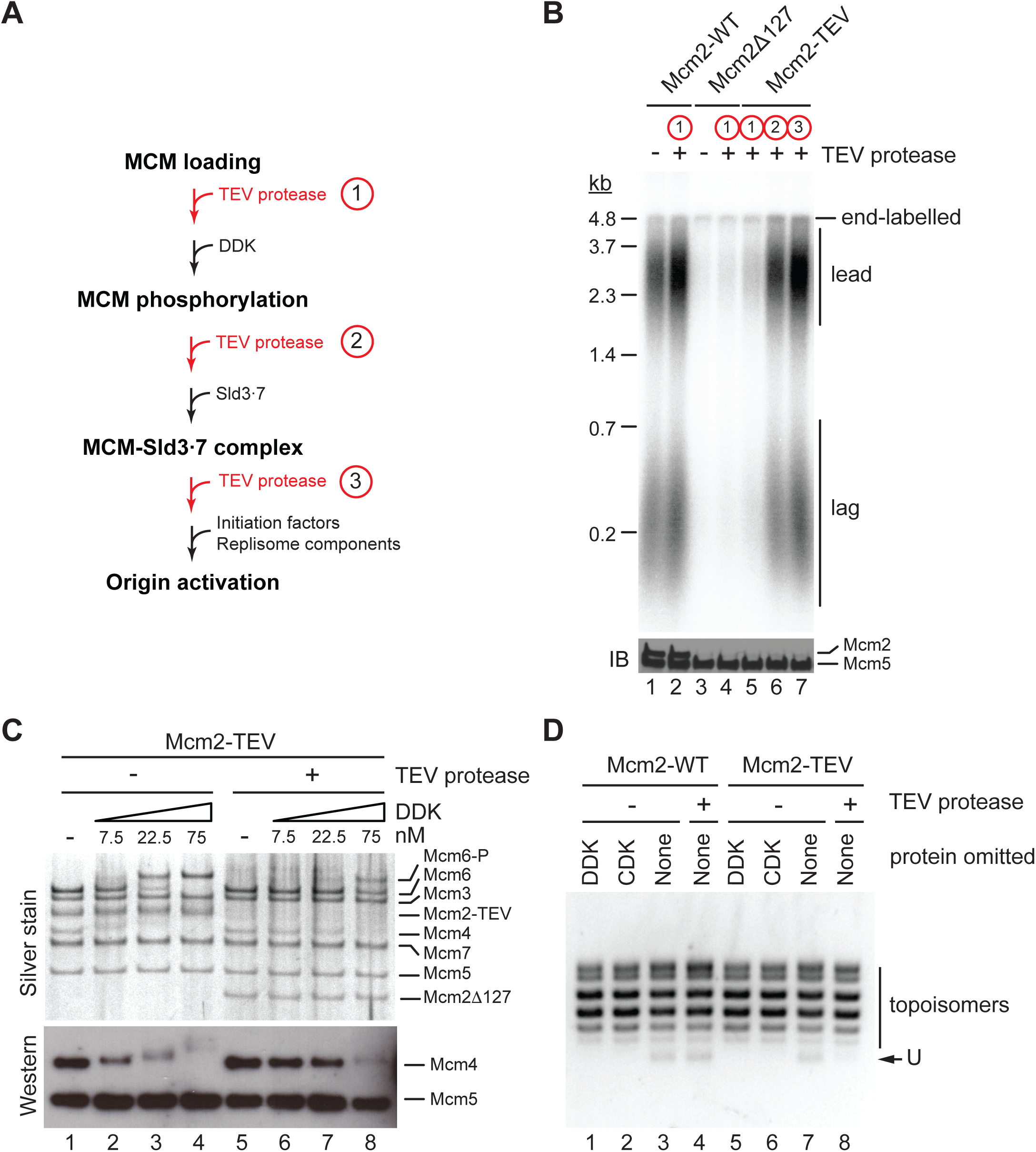
The Mcm2 N-terminal tail promotes DDK function during origin activation. (**A**) Experimental outline for experiment in B. Variable addition points for TEV protease are highlighted in red. (**B**) Standard *in vitro* DNA replication reactions were performed using p1017 (4.8 kb) as a template. TEV protease or mock buffer was added for 1 hour at 30°C as indicated. Reaction products were analyzed by denaturing agarose gel-electrophoresis and autoradiography (top). A fraction of each reaction was analyzed by SDS-PAGE and Western blot using antibodies against Mcm2 and Mcm5 (bottom); note that the N-terminal epitope recognized by the Mcm2 antibody is lost upon TEV protease cleavage. (**C**) Mcm2-7 DHs assembled with Cdt1·MCM^2-TEV^ were either mock-treated (lanes 1-4) or digested with TEV protease (lanes 5-8). DDK was subsequently added to the reactions at the indicated concentrations and reactions analyzed by SDS-PAGE and silver stain or Western blot using antibodies against Mcm4 and Mcm5. (**D**) Plasmid unwinding assay using CMGs assembled with either Cdt1·MCM^2-WT^ (lanes 1-4) or Cdt1·MCM^2-TEV^ (lanes 5-8) using p79 (3 kbp) as substrate. TEV protease was added to the reactions after the Mcm2-7 loading step, prior to the addition of DDK, CDK, Sld2, Sld3, Dpb11, GINS, Cdc45, Pol ε, RPA, and Mcm10 as indicated. DNA was repurified from the reaction and analyzed by native agarose gel-electrophoresis and EtBr stain. U: U-form DNA.

As the essential function of DDK is the phosphorylation of the N-terminal tails of Mcm4 and Mcm6, we asked whether proteolytic truncation of the Mcm2 N-terminal tail affects Mcm4 and -6 phosphorylation by DDK. For this we assembled Mcm2-7 DHs from Cdt1×MCM^2-TEV^ on bead-immobilized DNA and monitored DDK phosphorylation-dependent gel-mobility shifts of Mcm2-7 subunits by SDS-PAGE (**Figure 3C**). Mcm2-7 DHs harboring Mcm2-TEV were either digested or mock-treated with TEV protease prior to DDK addition. In the presence of the full-length Mcm2 N-terminal tail, Mcm4 and Mcm6, but not any of the other Mcm2-7 subunits, exhibited a pronounced retardation in gel-mobility in the presence of DDK. The disappearance of the unphosphorylated Mcm4 and -6 bands at higher DDK concentrations demonstrates that these subunits were phosphorylated quantitatively by DDK. In contrast, the DDK-dependent gel-retardation of Mcm4 and -6 was strongly diminished at all DDK concentrations tested when the Mcm2 N-terminal tail was truncated with TEV protease prior to DDK addition. Thus, the Mcm2 N-terminal tail promotes the phosphorylation of Mcm4 and Mcm6 by DDK in the context of Mcm2-7 DHs.

Phosphorylation of the Mcm4 and -6 N-terminal tails by DDK promotes the assembly of the CMG helicase. We, therefore, tested if truncation of the Mcm2 N-terminal tail impairs CMG assembly using an origin-dependent CMG helicase assay that detects CMG helicase activity by the generation of highly unwound circular plasmid DNA, termed U-form DNA (Douglas et al., 2018). As expected, generation of U-form DNA in the presence of either Mcm2-WT or Mcm2-TEV is dependent on both CDK and DDK, demonstrating that plasmid unwinding is dependent on CMG assembly (**Figure 3D**). Importantly, generation of U-form DNA was suppressed specifically in the presence of Mcm2-TEV when TEV protease was added to the reaction after the Mcm2-7 loading step, prior to the addition of DDK and other initiation factors. This data is consistent with residues 1-127 of the Mcm2 N-terminal tail promoting CMG assembly. In summary, we conclude that the Mcm2 N-terminal tail is important for DNA replication by promoting the phosphorylation of Mcm4 and -6 by DDK and subsequent CMG assembly, whereas it is dispensable for DNA replication after CMG assembly.

### The Mcm2 N-terminal tail promotes DDK docking onto Mcm2-7 DHs

Next we addressed how Mcm2 may promote the phosphorylation of Mcm4 and -6 by DDK. Previous studies had proposed a docking mechanism by which a stable association of DDK with Mcm2-7 DHs promotes processive multi-site phosphorylation of the Mcm4 and -6 N-terminal tails (Francis et al., 2009; Sheu and Stillman, 2006). These studies, however, either utilized complex cell extracts to load Mcm2-7 onto DNA, which may include unknown proteins that bridge the DDK-Mcm2-7 interaction, or examined the binding of DDK to the isolated Mcm4 subunit, leaving open the question how DDK interacts with MCM subunits in the context of the Mcm2-7 DH. We, therefore, investigated the direct binding of DDK to purified Mcm2-7 DHs *in vitro*. For this, Mcm2-7 DHs bound to DNA immobilized on paramagnetic beads were isolated from Mcm2-7 loading reactions, washed with high-salt buffer, and incubated under various conditions with purified DDK.

Titration of DDK into a Mcm2-7 DH binding reaction revealed that DDK binding to the DNA beads started to saturate at 75 nM DDK (**Figure 4A**). Importantly, when Mcm2-7 DH formation was prevented by omission of Cdt1×Mcm2-7 from the Mcm2-7 loading reaction DDK binding to the DNA beads was not observed even at 300 nM DDK, demonstrating that DDK binds specifically to Mcm2-7 DHs on the DNA. Next we tested the contribution of the Mcm2 N-terminal tail to the DDK-Mcm2-7 DH interaction. To this end, Mcm2-7 DHs harboring MCM2-TEV were incubated with TEV protease prior to addition of DDK. As before, proteolytic removal of residues 1-127 from the Mcm2 N-terminus did not affect Mcm2-7 DH stability on the DNA (**Figure 4B**). However, truncation of the Mcm2 N-terminus resulted in severely diminished binding of DDK to Mcm2-7 DHs, demonstrating that Mcm2 residues 1-127 are important for the association of DDK with Mcm2-7 DHs. To determine if the Mcm2 N-terminus is important only for the recruitment of DDK or also for the retention of DDK on Mcm2-7 DHs, TEV protease was added to the reaction either before or after DDK binding. As shown in **Figure 4C**, while truncation of the Mcm2 N-terminal tail prior to the addition of DDK inhibited DDK-Mcm2-7 DH complex formation as before, addition of TEV protease after DDK-Mcm2-7 DH complex formation also led to the release of DDK from Mcm2-7 DHs, demonstrating that the Mcm2 N-terminal tail is required for the retention of DDK on Mcm2-7 DHs.

**Figure 4.**
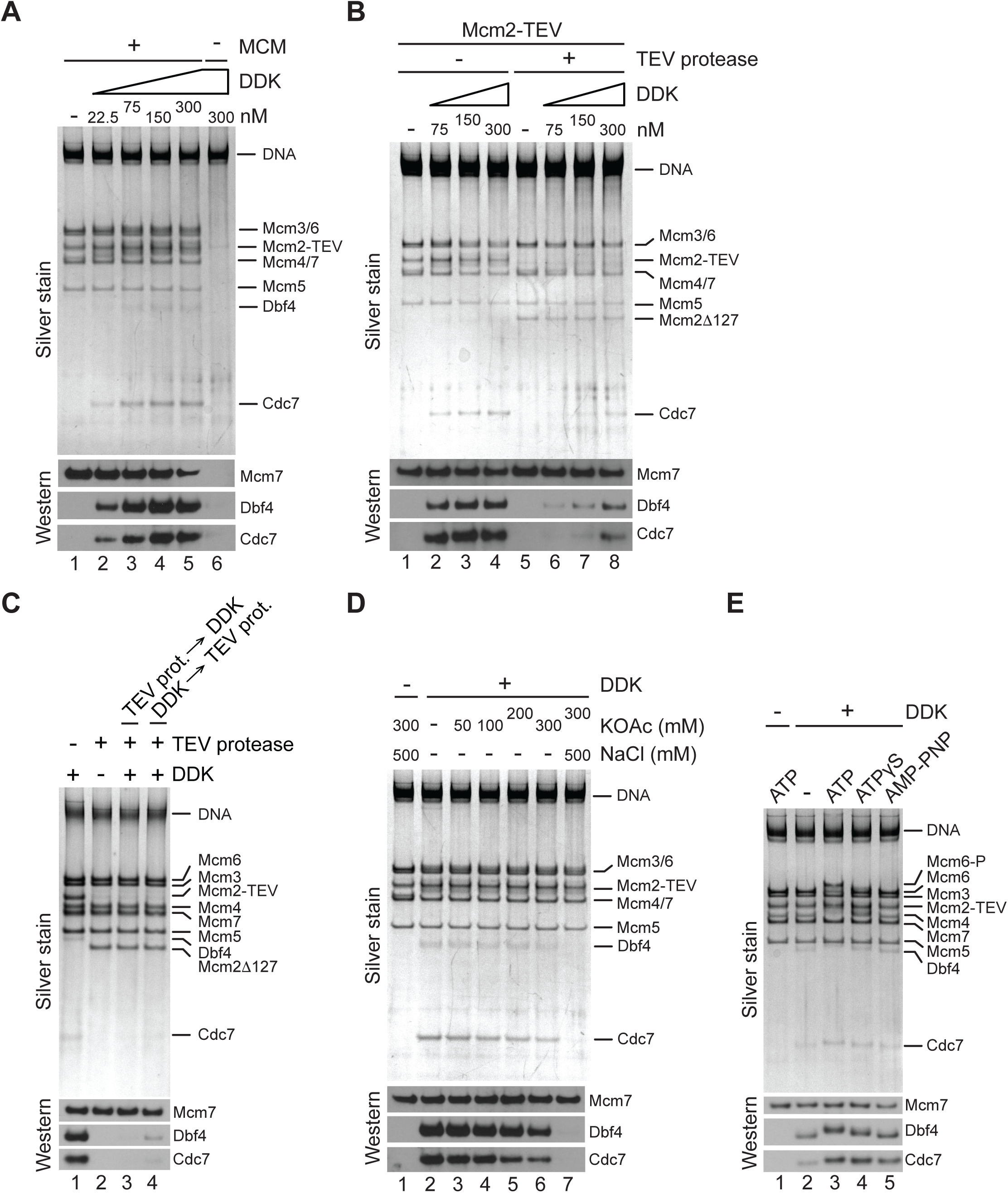
The Mcm2 N-terminal tail promotes binding of DDK to Mcm2-7 DHs. (**A**) Mcm2-7 DHs were assembled on bead-immobilized DNA, washed with high-salt buffer, and subsequently incubated with ATPγS and DDK at the indicated concentrations. As a control, Cdt1·Mcm2-7 was omitted from the Mcm2-7 loading reaction in lane 6. After incubation with DDK, DNA bound material was isolated and analyzed by SDS-PAGE and silver stain (top) or Western blot using antibodies against Mcm7, Dbf4, or Cdc7 (bottom). (**B**) Mcm2-7 DHs were assembled from Cdt1·MCM^2-TEV^, mock-treated or digested with TEV protease as indicated and incubated with purified DDK at the indicated concentrations. DNA-bound material was analyzed as in A. (**C**) Mcm2-7 DHs were assembled from Cdt1·MCM^2-TEV^ and mock-treated or digested with TEV protease as indicated. In lane 3, DDK was added after TEV protease, in lane 4 DDK was added before TEV protease. DNA-bound material was analyzed as in A. (**D**) DNA-bound DDK-Mcm2-7 DH complexes were washed with buffer containing the indicated concentration of KOAc, and where indicatewd followed by a wash with buffer containing 500 mM NaCl. (**E**) Mcm2-7 DHs were assembled on bead-immobilized DNA, washed to remove free ATP, and subsequently incubated with DDK in the presence of ATP or ATP analogues, as indicated. DNA-bound material was analyzed as in A.

The DDK-Mcm2-7 DH complex was remarkably resistant to extensive buffer washes including up to 0.3 M KOAc, attesting to the relative stability of the DDK-Mcm2-7 DH interaction (**Figure 4D**). Disruption of the complex was observed in wash buffer containing 0.5 M NaCl, a condition in which Mcm2-7 DHs remain stably bound to DNA, indicating that electrostatic interactions play an important role in mediating the DDK-Mcm2-7 DH interaction. In the above experiments DDK binding to Mcm2-7 DHs was monitored in the presence of the non-hydrolyzable ATP analogue ATPγS with the intention to trap DDK bound to Mcm2-7 DH. Indeed, reduced DDK binding to Mcm2-7 DHs is observed in the complete absence of ATP (**Figure 4E**). In contrast, the level of DDK binding to Mcm2-7 DHs was essentially identical in the presence of ATP, ATPγS, or the non-hydrolyzable ATP analogue adenylyl-imidodiphosphate (AMP-PNP). This demonstrates that DDK docking onto Mcm2-7 DHs is independent of Mcm4 and -6 phosphorylation. Moreover, these observations reveal that DDK remains stably bound to the Mcm2-7 DHs after phosphorylation of the Mcm4/-6 N-terminal tails. Together, these results demonstrate that the Mcm2 N-terminal tail is required for the physical interaction of DDK with Mcm2-7 DHs, explaining the loss of Mcm4 and -6 phosphorylation in its absence.

### Efficient origin activation requires the Mcm2 N-termini of both Mcm2-7 hexamers

The stoichiometry and molecular configuration of DDK bound to the Mcm2-7 DH during origin activation is not known. The symmetry of the Mcm2-7 double hexamer implies a complex of two DDK molecules per Mcm2-7 DH, with one DDK molecule bound to each hexamer. Alternatively, it may also be possible that one DDK molecule bound to the flexible Mcm2 N-terminus of one hexamer is sufficient to efficiently phosphorylate Mcm4 and -6 in both hexamers. To begin to address the mechanism of Mcm2-7 DH phosphorylation by DDK we assembled mixed Mcm2-7 DHs from wildtype Cdt1·MCM and Cdt1·MCM^2Δ127^. If both Mcm2 N-termini were required for origin activation, a 1:1 mixture of wildtype Cdt1·MCM and Cdt1·MCM^2Δ127^, which yields a mixed DH population composed of 25 % WT/WT, 50 % WT/Δ127, and 25 % Δ127/ Δ127 with respect to Mcm2, would be expected to cause a 75 % activity loss within a population of replication origins relative to wildtype Mcm2-7 DHs, whereas only a 25 % reduction would be expected if the Mcm2 N-terminus of one hexamer were sufficient to activate both hexamers.

To generate Cdt1·MCM^2Δ127^, we digested Cdt1·MCM^2-TEV^ with TEV protease and re-purified Cdt1·MCM^2Δ127^ from the digestion reaction by gel-filtration chromatography. Truncation of the Mcm2 N-terminus did not have a detectable effect on Cdt1·MCM stability, as Cdt1·MCM^2Δ127^ eluted in a mono-disperse peak during gel-filtration (**Figure 5A**). Cdt1·MCM^2Δ127^ also did not exhibit any discernible Mcm2-7 loading defects, demonstrating that the Mcm2 N-terminus is dispensable for Mcm2-7 recruitment and loading onto DNA (**Figure 5B**). Next we assessed the contribution of the Mcm2 N-terminus to DNA replication. For this, wildtype or mutant Cdt1·MCM complexes were included at 80 nM during the Mcm2-7 loading step, a concentration that supports near maximal DNA replication levels. Importantly, at Cdt1·MCM concentrations below 80 nM origin firing efficiency strongly correlates with Cdt1·MCM concentration, thus allowing sensitive detection of loss of Cdt1·MCM activity (**Supplementary Figure 4A**). Similar to our previous approach (**Figure 2**), total DNA synthesis in the complete absence of the Mcm2 residues 1-127 was reduced by ∼ 75 % relative to that in the presence of full-length Mcm2 (**Figures 5C+D**). This DNA synthesis defect was due to a defect in origin activation and not fork progression as leading and lagging strand lengths were similar in both conditions (**Figure 5E**). Importantly, DNA replication was reduced by ∼ 50 % at a 1:1 ratio of Cdt1·MCM^2Δ127^ to wildtype Cdt1·MCM, well beyond the 25 % reduction expected if one Mcm2 N-terminal tail was sufficient for normal origin activation. The fact that the loss in DNA replication levels falls short of the 75 % reduction expected if both Mcm2 N-terminal tails in the Mcm2-7 DH were required for origin firing is attributable to the residual origin activity observed in the absence of Mcm2 N-terminal tails (**Supplementary Figure 4B**). Importantly, we did not detect any evidence for asymmetric, or uni-directional, origin firing arising from the activation of a single hexamer within heterologous Mcm2-7 DHs assembled from both Cdt1×MCM^2-WT^ or Cdt1×MCM^2Δ127^, which may either generate greater than half-unit length leading strands, or cause a disproportional increase in lagging strand products as leading strand synthesis is initiated by the lagging strand of the sister replisome (Aria and Yeeles, 2018). In summary, we conclude that the Mcm2 N-termini of both hexamers are required for efficient origin firing, while the presence of a single Mcm2 N-terminal tail at a Mcm2-7 DH does not result in the establishment of single replication forks at an origin.

**Figure 5.**
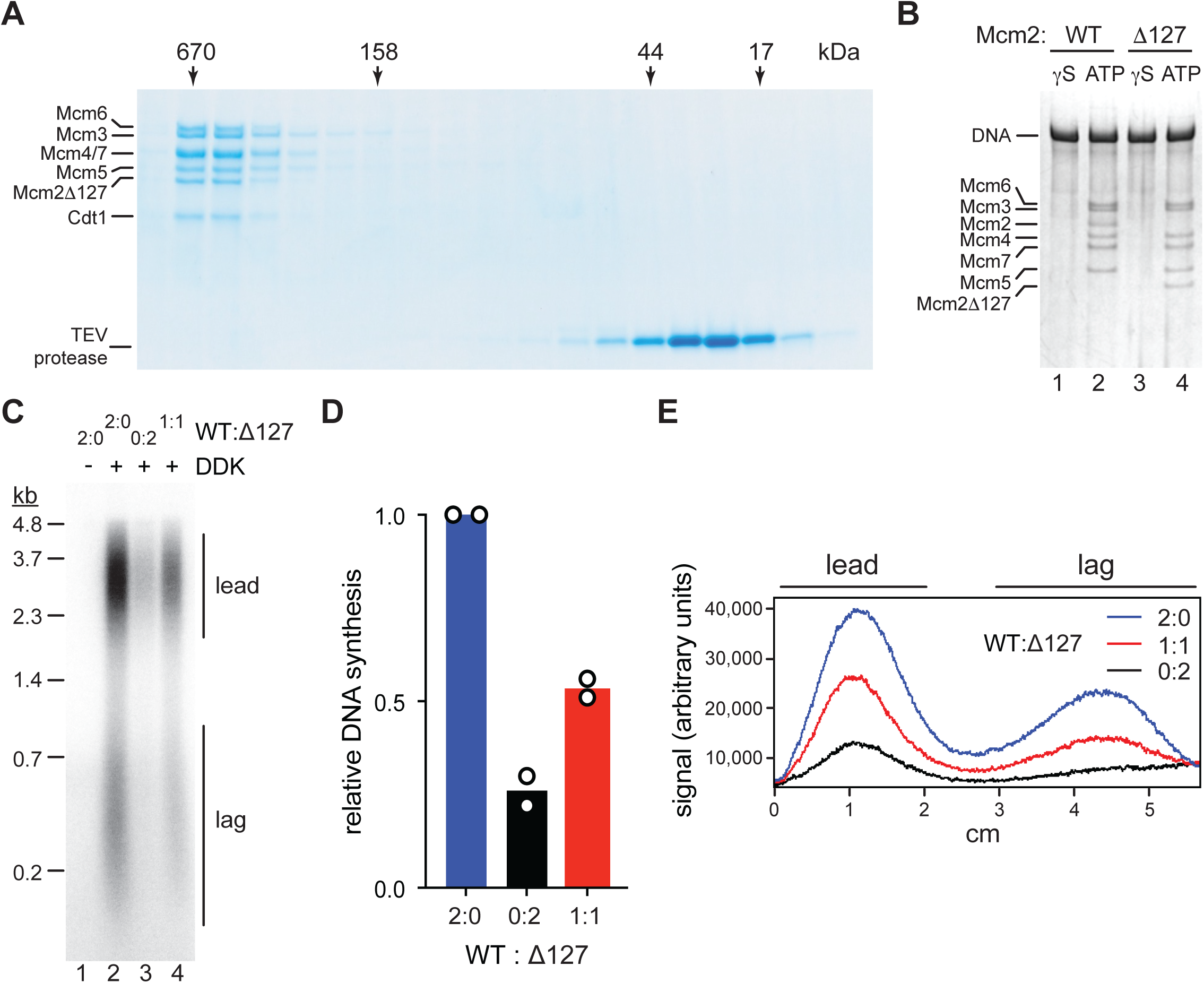
Mcm2-WT is not able to rescue Mcm2Δ127 replication defect. (**A**) Gel-filtration analysis of purified Cdt1·MCM^2-TEV^ following digestion with TEV protease. The digestion reaction was fractionated on a Superdex 200 column and fractions analyzed by SDS-PAGE and Coomassie stain. (**B**) Mcm2-7 loading reactions with either wildtype Cdt1·MCM (lanes 1+2) or Cdt1·MCM^2-Δ127^ (lanes 3+4). Reactions were performed either in the presence of ATPγS or ATP as indicated and DNA-beads subsequently washed with high-salt buffer. DNA-bound fraction was analyzed by SDS-PAGE and silver stain. (**C**) Standard DNA replication reaction using p1017 (4.8 kb) as template. Cdt1·MCM^2-Δ127^ and Cdt1·MCM^2-WT^ were included at the Mcm2-7 loading step at the indicated ratios; the total concentration of Cdt1·MCM was 80 nM in the Mcm2-7 loading reaction. (**D**) Quantification of total relative DNA synthesis in reactions of experiment in C. Bars represent the average of two independent experiments. (**E**) Lane traces of experiment in C.

### Rad53 sterically inhibits DDK binding to Mcm2-7 DHs

We have shown that DDK binding to Mcm2-7 DHs is required for origin activation. Interestingly, previous studies have indicated that a physical interaction between Rad53 and DDK contributes to the inhibition of origin activation by the checkpoint (Chen et al., 2013; Duncker et al., 2002; Matthews et al., 2014; Varrin et al., 2005). We, therefore, asked whether Rad53 might control origin activity by inhibiting DDK binding to Mcm2-7 DHs. For this we purified recombinant wildtype Rad53, which undergoes autoactivation during overexpression in *E. coli*, or the catalytically dead Rad53-D339A mutant, designated Rad53-kd below (Gilbert et al., 2001). Indeed, pre-incubation of DDK with Rad53 in the presence of ATP prior to addition of Mcm2-7 DHs prevented both the binding of DDK to Mcm2-7 DHs and Mcm4 and -6 phosphorylation by DDK (**Figure 6A**, lanes 4+5). Moreover, Rad53 was able to displace DDK from Mcm2-7 DHs when DDK binding to Mcm2-7 DHs preceded addition of Rad53 (lanes 4+7). However, this displacement action of Rad53 was slightly less efficient at disrupting the DDK-Mcm2-7 DH interaction than the action of preventing DDK recruitment (lanes 5+7). Intriguingly, Rad53-kd also largely inhibited stable binding of DDK to Mcm2-7 DHs when co-incubated with DDK prior to addition to Mcm2-7 DHs, demonstrating that DDK phosphorylation by Rad53 is not essential to prevent stable DDK recruitment to Mcm2-7 (lanes 4+6). However, some residual DDK binding and significant Mcm4 and -6 phosphorylation occurred in the presence of Rad53-kd, indicating that Rad53-kd-mediated inhibition of DDK is inefficient. Moreover, unlike Rad53-WT, Rad53-kd was unable to displace DDK from Mcm2-7 DHs when DDK was bound to Mcm2-7 DHs prior to Rad53-kd addition (lanes 4+8). These observations are consistent with genetic data demonstrating that kinase-dead alleles of Rad53 are deficient in origin inhibition (Lopes et al., 2001; Pellicioli et al., 1999). We conclude that activated Rad53 inhibits DDK binding to Mcm2-7 DHs, which in turn inhibits Mcm4 and -6 phosphorylation.

**Figure 6.**
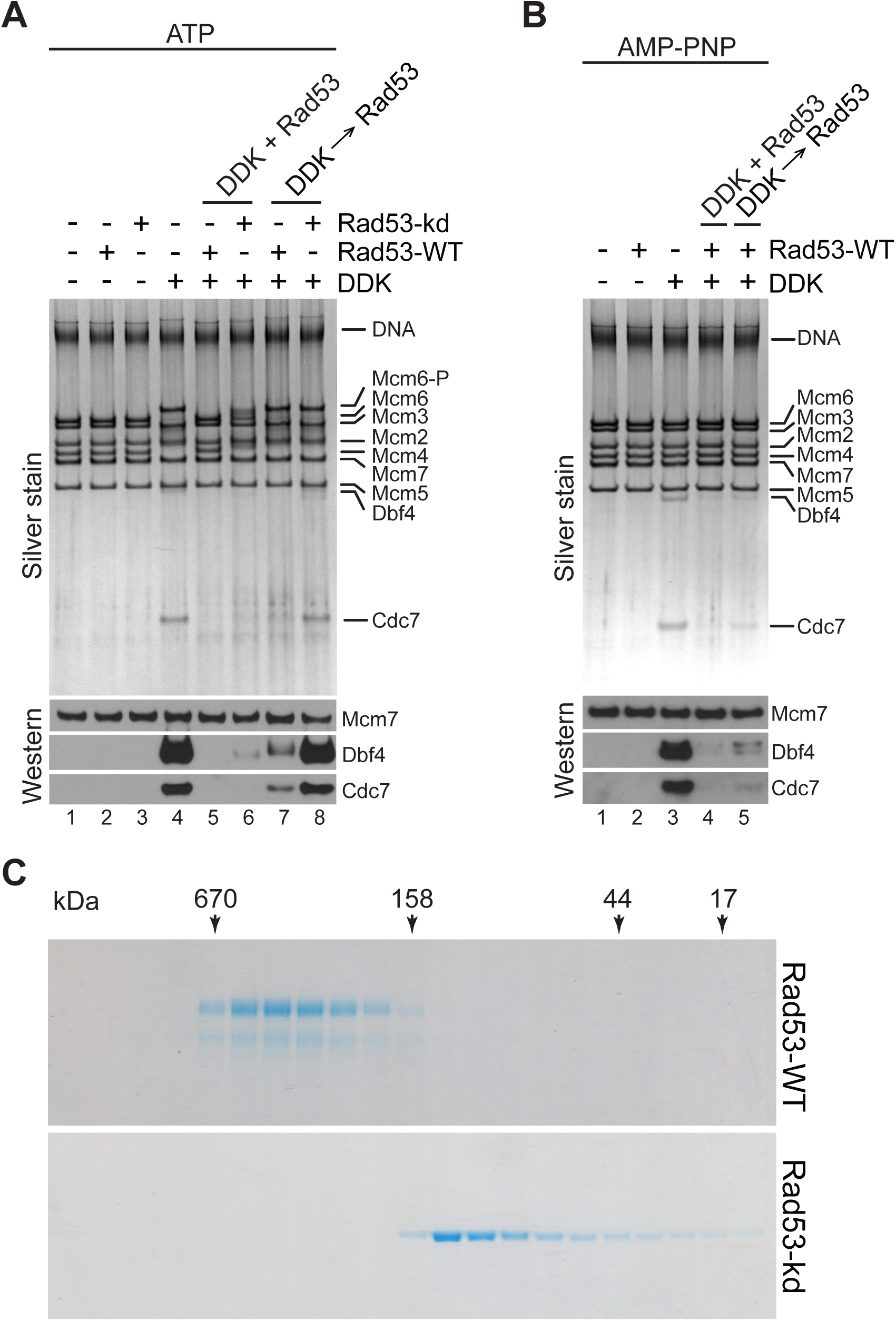
Steric inhibition of DDK binding to Mcm2-7 DHs by Rad53. (**A**) DDK binding to purified Mcm2-7 DHs was monitored in the presence of ATP and in the absence or presence of Rad53-WT or Rad53-kd, as indicated. In lanes 5+6 DDK and Rad53 were co-incubated in the presence of ATP prior to addition to DNA-bound Mcm2-7 DHs; in lanes 7+8 DDK was incubated with purified Mcm2-7 DHs before addition of Rad53. DNA-bound material was analyzed SDS-PAGE and silver stain or Western blot as indicated. (**B**) DDK binding to DNA-bound Mcm2-7 DHs was monitored in the presence of AMP-PNP. DDK and Rad53 were either co-incubated in the presence of AMP-PNP prior to addition to purified DNA-bound Mcm2-7 DHs (lane 4), or added sequentially to Mcm2-7 DHs (lane 5) as indicated. (**C**) Gel filtration profiles of purified Rad53-WT and Rad53- kd. Fractions were analyzed by SDS-PAGE and Coomassie stain.

To further address if Rad53 phosphorylation of DDK is required for the inhibition of DDK binding to Mcm2-7 DHs we monitored DDK-Mcm2-7 DH complex formation in the presence of AMP-PNP. As we have shown above, AMP-PNP promotes DDK binding to Mcm2-7 DHs to the same extent as ATP (**Figure 4E**). Strikingly, the ability of Rad53 to prevent DDK recruitment to Mcm2-7 DHs or to displace DDK from Mcm2-7 DHs was undiminished in the presence of AMP-PNP (**Figure 6B**). Thus, DDK phosphorylation by Rad53 is not required to control DDK-Mcm2-7 DH complex formation. This raises the question why Rad53-kd is deficient at inhibiting DDK. Intriguingly, Rad53-WT and Rad53-kd exhibit very different elution profiles during gel-filtration chromatography (**Figure 6C**). While Rad53-kd elutes at a volume that is consistent with a monomeric structure, Rad53-WT elutes as oligomeric complex. Rad53 and its human homolog, Chk2, are known to associate into homo-dimeric complexes during activation by trans-autophosphorylation, suggesting the oligomeric form of Rad53 observed here is a dimer (Ahn and Prives, 2002; Cai et al., 2009; Oliver et al., 2006; Wybenga-Groot et al., 2014; Xu et al., 2002). However, contrary to our observation, both kinase-dead Rad53 and Chk2 have been shown previously to also form dimers (Cai et al., 2009; Wybenga-Groot et al., 2014). As these previous studies were carried out with truncated kinase versions it is possible that the additional domains present in our full-length Rad53-kd construct affect its dimerization. For example, the isolated Chk2 kinase domain adopts a highly distinct dimer configuration from that of a Chk2 construct spanning the FHA and kinase domain, supporting the notion that domain composition can affect the oligomeric structure of Rad53/Chk2 (Cai et al., 2009; Oliver et al., 2006). We conclude that Rad53 activation results in the formation of Rad53 dimers that can sterically inhibit DDK binding to Mcm2-7 DHs, suggesting a novel non-catalytic mechanism for Rad53- dependent origin control.

## Discussion

The head-to-head orientation of the hexamers in the Mcm2-7 DH requires CMG helicases to pass each other during origin firing (Douglas et al., 2018; Georgescu et al., 2017). Based on this configuration it was proposed that an inactive CMG encircling dsDNA blocks the progression of a CMG formed around the opposite hexamer to impose bidirectional origin firing (Douglas et al., 2018; Georgescu et al., 2017). However, a stalled CMG helicase encircling ssDNA would pose a threat to DNA integrity due to the exposure of the displaced strand at the active CMG. As an alternative fail-safe mechanism we propose that simultaneous activation of both Mcm2-7 hexamers at an origin is controlled by an interdependent mechanism. Our data identify the Mcm2 N-terminal tail as a critical component of such a mechanism, as loss of a single Mcm2 N-terminus at a Mcm2-7 DH inhibits origin activity without inducing unidirectional firing. Mechanistically, we propose that both Mcm2 N-terminal tails in a Mcm2-7 DH are required for tethering of a pair of DDK molecules to promote symmetric phosphorylation of both hexamers (**Figure 7A**). Supporting an interdependent hexamer activation mechanism, recent findings have demonstrated that both Mcm2-7 hexamers loaded at an origin have to be physically associated to support CMG activation (Champasa et al., 2019). Interdependent Mcm2-7 activation may be required at the origin melting step, which may involve two CMGs working in opposite direction against each other (Froelich et al., 2014; Langston and O’Donnell, 2019; Noguchi et al., 2017).

**Figure 7.**
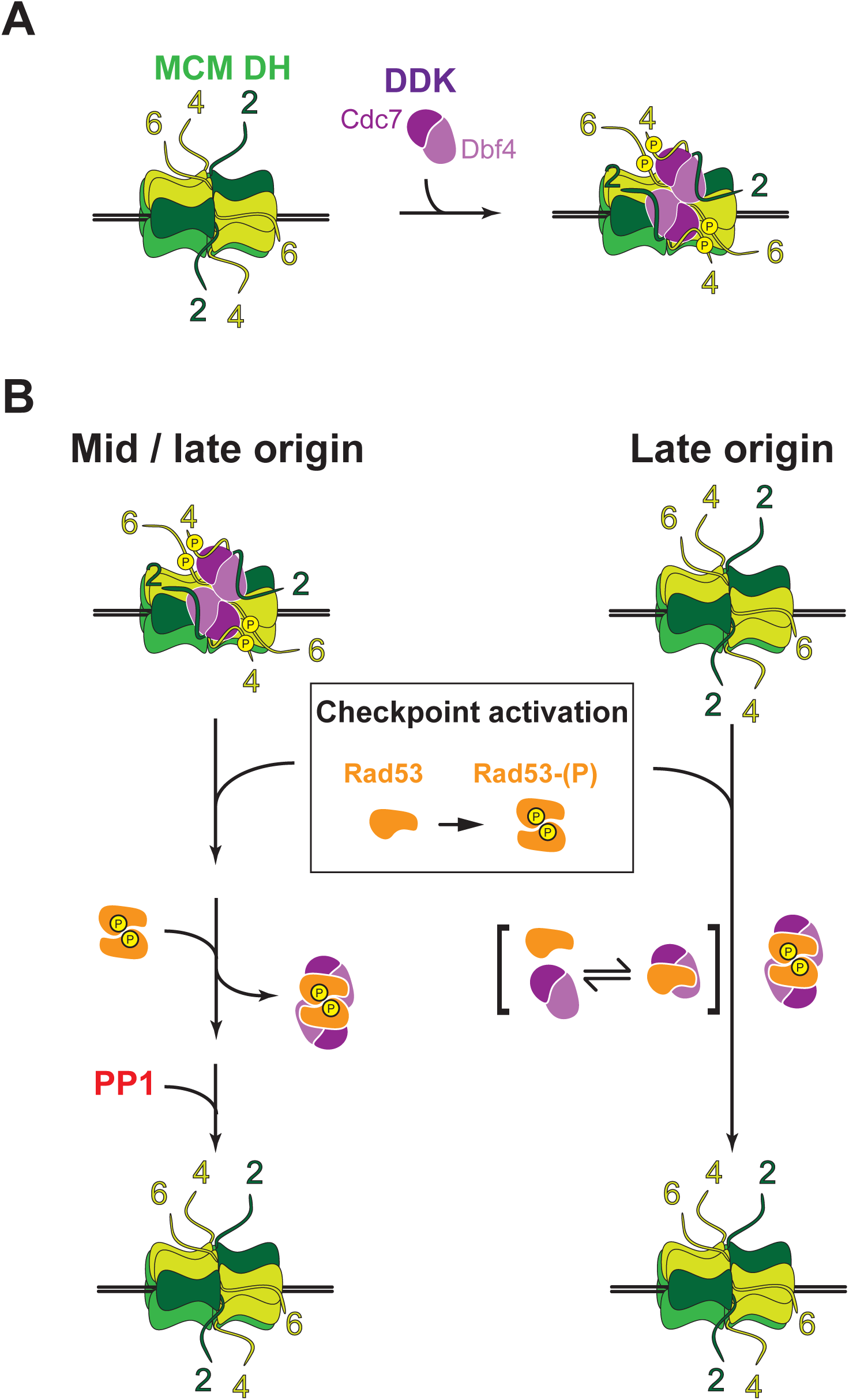
Model for the recruitment of DDK to Mcm2-7 DHs and its regulation Rad53. (**A**) Two DDK molecules are recruited to a Mcm2-7 DH via the Mcm2 N-terminal tails. Upon recruitment to the Mcm2-7 DH, DDK phosphorylates the Mcm4 and -6 N-terminal tails. Following Mcm4 and -6 phosphorylation both DDK molecules remain stably associated with the Mcm2-7 DH via the Mcm2 N-terminal tails, possibly in the form of a DDK dimer. Numbers denote the MCM subunit identity of the respective flexible N-terminal tails. (**B**) Checkpoint activation after early origin firing in S phase induces the dimerization and trans-autophosphorylation of Rad53. At origins that have already recruited DDK but have not yet fired, activated Rad53 dimers displace DDK from the Mcm2-7 DH, which allows PP1 to dephosphorylate and inactivate the Mcm2-7 DH (left). Active Rad53 dimers also prevent the recruitment of soluble DDK to late origins (right). Monomeric inactive Rad53 can form unstable complexes with soluble DDK (bracketed) that can impair DDK recruitment to Mcm2-7 DHs, but not completely block Mcm4/6 phosphorylation by DDK.

Our proposed role of the Mcm2 N-terminal tail as a DDK docking site is supported by a previous yeast two-hybrid study that detected a specific pairwise interaction between Dbf4 and the Mcm2 N-terminus (Ramer et al., 2013). Although a docking interaction between DDK and the structured NTD of Mcm4 was suggested by *in vitro* studies with purified DDK and Mcm4 (Sheu and Stillman, 2006), we find that in the context of the Mcm2-7 DH this interaction is insufficient to tether DDK in the absence of the Mcm2 N-terminal tail. However, our data does not rule out the possibility that DDK docking involves a complex interaction surface comprising critical interactions with multiple Mcm2-7 subunits. In fact, since Mcm2-7 DHs are preferred targets for DDK over single Mcm2-7 hexamers additional structural determinants are likely to govern the interaction of DDK with Mcm2-7 (Francis et al., 2009; Sun et al., 2014). In addition, prior phosphorylation of Mcm2-7 by other kinases has also been shown to promote DDK binding to Mcm2-7 at origins (Francis et al., 2009). Structural characterization of the DDK·Mcm2-7 DH complexes isolated here will help define the details of the DDK- Mcm2-7 interface. Such an analysis may also help resolve the mechanism of Mcm4/6 phosphorylation by DDK, which may occur across the hexamer-hexamer interface or within the hexamer bound by a DDK molecule (Sun et al., 2014).

We find that DDK remains stably bound to Mcm2-7 DHs even after phosphorylation of the Mcm4 and -6 N-terminal tails. DDK phosphorylation of Mcm2-7 is opposed by dephosphorylation of Mcm2-7 by the Rif1-PP1 phosphatase complex (Alver et al., 2017; Dave et al., 2014; Hiraga et al., 2014; Hiraga et al., 2017; Mattarocci et al., 2014). While tethering of DDK to Mcm2-7 DHs may, therefore, not be essential for maintaining DDK phosphorylation of Mcm4/6 in the absence of Rif1-PP1 *in vitro* (**Figure 3B**, lane 6 and **Figure 4C**, lane 4), it is likely to promote Mcm2-7 phosphorylation *in vivo* by shifting the balance between DDK and Rif1-PP1 activity towards DDK. Additionally, as Dbf4 concentrations are limiting for origin activation, retention of DDK at early origins until origin firing may prevent premature recycling of DDK to late origins to establish the replication timing program (Mantiero et al., 2011; Tanaka et al., 2011). Similarly, retention of DDK at unfired origins has been proposed to restrict DDK targeting of Eco1 until late S phase to control cohesion in the cell cycle (Seoane and Morgan, 2017). These mechanisms imply that DDK release from Mcm2-7 DHs is regulated by origin activation. Since DDK targets Mcm2-7 DHs specifically, hexamer separation may be sufficient to induce DDK release (Francis et al., 2009; Sun et al., 2014). Alternatively, tethering of DDK to the long and flexible Mcm2 N-terminal tail may allow DDK to remain bound to replisomes. Indeed, the association of DDK with replisomes was shown to link DNA replication with meiotic recombination (Murakami and Keeney, 2014). Analogously, DDK bound to replisomes in a mitotic S phase may promote spatio-temporal coordination of DNA replication with other chromosomal processes such as chromatin assembly or DNA repair (Furuya et al., 2010; Gerard et al., 2006). It will, therefore, be interesting in future experiments to determine the dynamics of the DDK-Mcm2-7 interaction during DNA replication.

We find that activated Rad53 inhibits DDK binding to Mcm2-7 DHs by a steric mechanism that is independent of Dbf4 phosphorylation by Rad53. Dbf4 contains three evolutionarily conserved sequence motifs, termed N, M, and C that constitute parts of distinct domains separated by flexible linkers (Hughes et al., 2012; Masai and Arai, 2000). The M and C domains embrace the Cdc7 kinase to mediate Cdc7 activation, while the N motif forms part of an N-terminal BRCT domain that is dispensable for Cdc7 activation (Almawi et al., 2016; Hughes et al., 2012). Intriguingly, an N-terminal fragment encompassing the BRCT and M domains is required and sufficient to target Dbf4 to Mcm2-7 complexes loaded at replication origins (Dowell et al., 1994; Francis et al., 2009), while a physical interaction between Rad53 and the Dbf4 BRCT domain has been proposed to contribute to checkpoint-dependent origin control (Chen et al., 2013; Duncker et al., 2002; Matthews et al., 2012; Matthews et al., 2014). Thus, competitive binding of Rad53 to the Dbf4 BRCT domain is likely in part responsible for the inhibition of DDK binding to Mcm2-7 DHs observed here. The N-terminal region of Dbf4 has also been shown to interact with ORC (Duncker et al., 2002). However, consistent with a previous study in whole-cell yeast extracts (Francis et al., 2009), we show here that purified Mcm2-7 DHs can recruit DDK to replication origins independently of ORC.

Both Rad53-WT and Rad53-kd can inhibit DDK recruitment to Mcm2-7 DHs, but the block of Mcm4 and -6 phosphorylation in the presence of Rad53-kd is incomplete. More strikingly, unlike Rad53-WT, Rad53-kd is completely deficient for displacing DDK from Mcm2-7 DHs. However, as Rad53-dependent DDK displacement from Mcm2-7 DHs does not require Rad53 kinase activity, we suggest that the oligomeric state of Rad53, determined by the trans-autophosphorylation activity of Rad53, controls the ability of Rad53 to displace DDK from Mcm2-7 DHs. While Rad53-kd is monomeric in solution, we show that active Rad53-wt forms oligomers, likely dimers, in solution. We propose that following recruitment to Mcm2-7 DHs, DDK may form a dimer on the Mcm2-7 DH surface that is in complex with both Mcm2 N-terminal tails, which we show are required for maintaining DDK at Mcm2-7 DHs. A DDK dimer bound to a Mcm2-7 DH may require simultaneous disruption of the interaction of both DDK molecules with Mcm2-7 for DDK release, explaining the proficiency of dimeric Rad53-WT and, conversely, deficiency of monomeric Rad53-kd to compete DDK off Mcm2-7 DHs. Following DDK displacement, PP1 would then be able to dephosphorylate the Mcm4 and -6 N-terminal tails and thus inactivate the origin (**Figure 7B**, left). On the other hand, as DDK is monomeric in solution, both Rad53-WT and Rad53-kd can sequester DDK prior to Mcm2-7 DH binding, but the interaction of Rad53-kd with DDK is unstable (**Figure 7B**, right). Thus, Rad53 would block DDK binding to Mcm2-7 DHs *in vivo* only after checkpoint-induced activation and dimerization, as expected.

A Rad53 kinase-independent mechanism for DDK inhibition was unexpected as previous studies have demonstrated that mutation of Rad53 phosphorylation sites in Dbf4 and Sld3 allows late origin firing in the presence of HU or MMS (Lopez-Mosqueda et al., 2010; Zegerman and Diffley, 2010). It is possible that origin efficiency under this condition will further increase upon disruption of the Rad53-DDK interface. Alternatively, phosphoacceptor-site mutations in Dbf4 may also affect the physical interaction between Dbf4 and Rad53. These possibilities remain to be tested in the future.

## Materials and Methods

### Materials

Yeast strains, plasmids, oligos, and antibodies are listed in Tables 1-4.

**Table 1:**
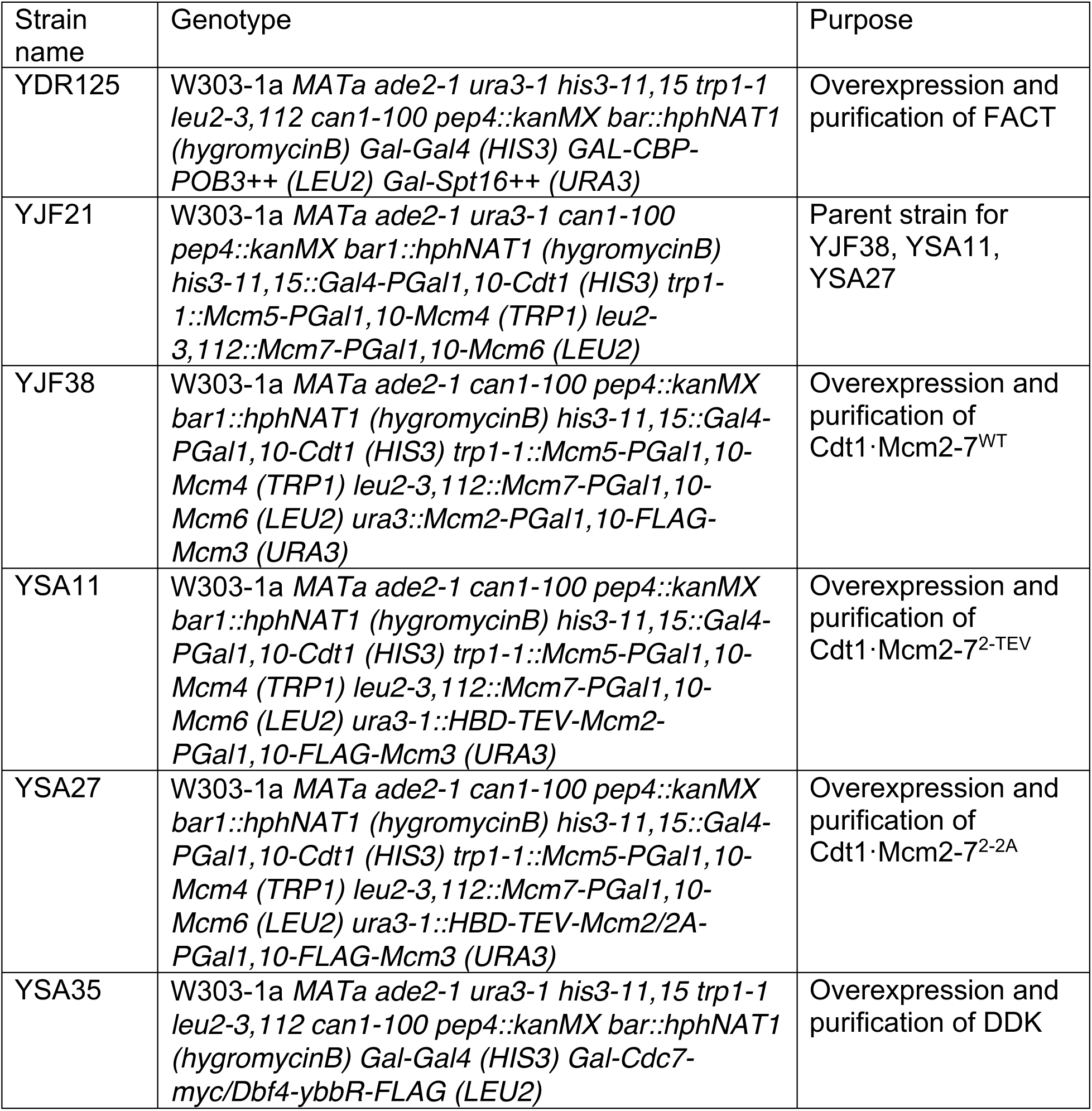
Yeast strains

| Strain name | Genotype | Purpose |
| --- | --- | --- |
| YDR125 | W303-1a *MATa ade2-1 ura3-1 his3-11,15 trp1-1 leu2-3,112 can1-100 pep4::kanMX bar::hphNAT1 (hygromycinB) Gal-Gal4 (HIS3) GAL-CBP-POB3++ (LEU2) Gal-Spt16++ (URA3)* | Overexpression and purification of FACT |
| YJF21 | W303-1a *MATa ade2-1 ura3-1 can1-100 pep4::kanMX bar1::hphNAT1 (hygromycinB) his3-11,15::Gal4-PGal1,10-Cdt1 (HIS3) trp1-1::Mcm5-PGal1,10-Mcm4 (TRP1) leu2-3,112::Mcm7-PGal1,10-Mcm6 (LEU2)* | Parent strain for YJF38, YSA11, YSA27 |
| YJF38 | W303-1a *MATa ade2-1 can1-100 pep4::kanMX bar1::hphNAT1 (hygromycinB) his3-11,15::Gal4-PGal1,10-Cdt1 (HIS3) trp1-1::Mcm5-PGal1,10-Mcm4 (TRP1) leu2-3,112::Mcm7-PGal1,10-Mcm6 (LEU2) ura3::Mcm2-PGal1,10-FLAG-Mcm3 (URA3)* | Overexpression and purification of Cdt1·Mcm2-7<sup>WT</sup> |
| YSA11 | W303-1a *MATa ade2-1 can1-100 pep4::kanMX bar1::hphNAT1 (hygromycinB) his3-11,15::Gal4-PGal1,10-Cdt1 (HIS3) trp1-1::Mcm5-PGal1,10-Mcm4 (TRP1) leu2-3,112::Mcm7-PGal1,10-Mcm6 (LEU2) ura3-1::HBD-TEV-Mcm2-PGal1,10-FLAG-Mcm3 (URA3)* | Overexpression and purification of Cdt1·Mcm2-7<sup>2-TEV</sup> |
| YSA27 | W303-1a *MATa ade2-1 can1-100 pep4::kanMX bar1::hphNAT1 (hygromycinB) his3-11,15::Gal4-PGal1,10-Cdt1 (HIS3) trp1-1::Mcm5-PGal1,10-Mcm4 (TRP1) leu2-3,112::Mcm7-PGal1,10-Mcm6 (LEU2) ura3-1::HBD-TEV-Mcm2/2A-PGal1,10-FLAG-Mcm3 (URA3)* | Overexpression and purification of Cdt1·Mcm2-7<sup>2-2A</sup> |
| YSA35 | W303-1a *MATa ade2-1 ura3-1 his3-11,15 trp1-1 leu2-3,112 can1-100 pep4::kanMX bar::hphNAT1 (hygromycinB) Gal-Gal4 (HIS3) Gal-Cdc7-myc/Dbf4-ybbR-FLAG (LEU2)* | Overexpression and purification of DDK |

**Table 2:** Plasmids

| Plasmid name | Description | Resistance marker | Auxotrophic marker | Purpose |
| --- | --- | --- | --- | --- |
| p79 | ARS1-containing pUC19 | Amp | - | Plasmid unwinding assay |
| p470 | ARS305-containing pUC19 derivative | Amp | - | Template for MCM loading, phosphorylation, DDK binding, and replication assays |
| p779 | pRS306G-MCM2/FLAG-MCM3 | Amp | URA3 | Yeast overexpression of Mcm2 and FLAG-Mcm3, template for Mcm2 modifications |
| p993 | pRS305G-CBP-POB3++ | Amp | LEU2 | Yeast overexpression of CBP-Pob3 |
| p1000 | pRS306G-SPT16++ | Amp | URA3 | Yeast overexpression of Spt16 |
| p1017 | ARS1-containing CEN4- pUC119 | Amp | - | Template for replication assay |
| p1034 | pRS306G-HBD-TEV-MCM2/FLAG-MCM3 | Amp | URA3 | Yeast overexpression of Mcm2-TEV and FLAG-Mcm3 |
| p1035 | pet15b-Nhp6 | Amp | - | Bacterial overexpression of 6xHis-Nhp6 |
| p1155 | pRS306G-Mcm2-Y82A/FLAG-Mcm3 | Amp | URA3 | Template for Mcm2 mutation |
| p1162 | pRS306G-Mcm2-2A/FLAG-Mcm3 | Amp | URA3 | Yeast overexpression of Mcm2-2A and FLAG-Mcm3 |
| p1220 | pRS305G-Cdc7-myc/Dbf4-ybbR-FLAG | Amp | LEU2 | Yeast overexpression of DDK with Cdc7-myc and Dbf4-ybbR-FLAG |

**Table 3:**
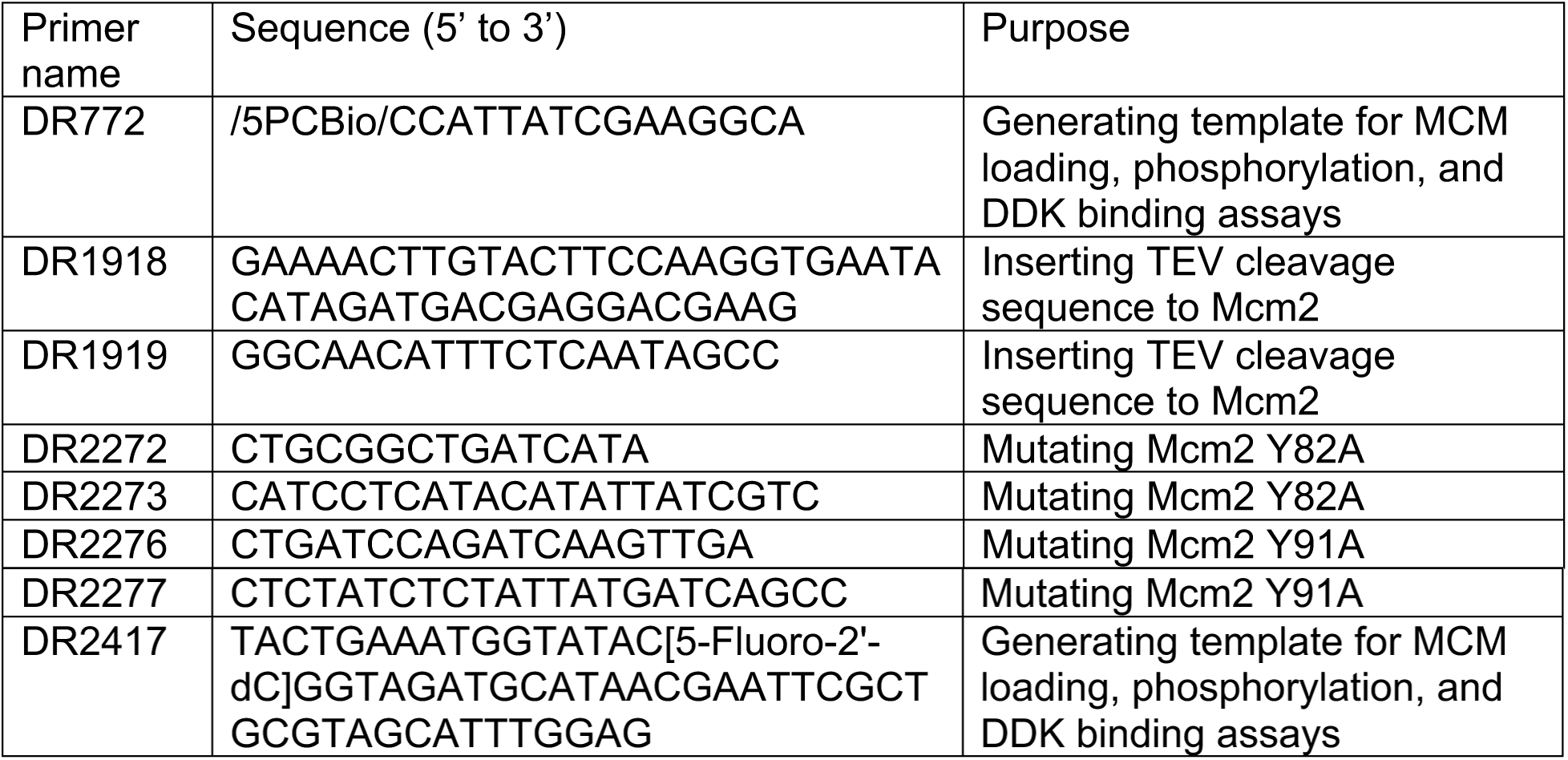
Oligos

**Table 4:** Antibodies

| Epitope | Company | Product number | Dilution |
| --- | --- | --- | --- |
| Cdc7 (yN-18) | Santa Cruz Biotechnology | sc-11964 | 1:2,000 |
| Dbf4 (yA-16) | Santa Cruz Biotechnology | sc-5706 | 1:2,000 |
| Mcm2 (yN-19) | Santa Cruz Biotechnology | sc-6680 | 1:2,000 |
| Mcm4 (yC-19) | Santa Cruz Biotechnology | sc-6685 | 1:1,000 |
| Mcm5 (yN-19) | Santa Cruz Biotechnology | sc-6687 | 1:2,000 |
| Mcm7 (yN-19) | Santa Cruz Biotechnology | sc-6688 | 1:2,000 |
| Goat IgG-HRP | Santa Cruz Biotechnology | sc-2354 | 1:5,000 |

### Protein purification

Proteins were purified as described previously, unless specified below (Devbhandari et al., 2017; Devbhandari and Remus, 2020).

#### DDK

Two DDK variants, harboring either a removable C-terminal TAP^tcp^ tag (Gros et al., 2014) or a ybbR-FLAG tag on Dbf4 were used interchangeably. Both variants behave identically and are fully proficient for DNA replication *in vitro*.

The ybbR-FLAG-tagged DDK was purified from strain YSA35. Cells were grown in 48L YP / 2 % glycerol / 2 % lactic acid pH 5.5 (YPLG) at 30°C to a density of 2 × 10^7^ cells / mL. Protein expression was induced with 2 % galactose for 4 hours. Cells were collected by centrifugation, washed with 25 mM HEPES-KOH pH 7.6 / 1 M sorbitol and resuspended in 0.5 volumes of buffer A (45 mM HEPES-KOH pH 7.6 / 0.02 % NP-40 substitute / 10 % glycerol) / 100 mM NaCl / 1mM DTT / 1x protease inhibitor cocktail (Pierce). The cell suspension was pipetted dropwise into liquid nitrogen to generate frozen popcorn and stored in −80°C. Cells were lysed by crushing the popcorn in a Spex freezer mill, using 10 cycles of 2 minute run + 1 minute cooldown at 15 CPS. Resulting whole cell lysate was thawed and supplemented with 1 volume of 45 mM buffer A / 100mM NaCl / 1mM DTT / 1x protease inhibitor cocktail. 5 M NaCl was added to the lysate to a final concentration to 300 mM. After 20 minutes of gentle agitation at 4°C, cell lysate was centrifuged in a T-647.5 rotor (Thermo Fisher) at 40,000 rpm for 1 hour at 4°C. The clear soluble phase was recovered and DDK pulled down with 1 mL packed FLAG affinity agarose beads (Sigma) for 4 hours at 4°C with gentle rocking. Beads were collected by centrifugation and washed with 10 volumes of buffer A / 300 mM NaCl/1 mM DTT. Beads were resuspended in 1 volume of buffer A / 300 mM NaCl / 2 mM MnCl_2_ / 1 mM DTT and incubated with λ protein phosphatase (NEB) at 50 U / mL for 1 hour at 23°C with agitation. Beads were collected and protein eluted in 5 volumes of buffer A / 300 mM NaCl / 1 mM DTT supplemented with 0.25 mg / mL 3xFLAG peptide. Eluates were analyzed by SDS-PAGE. The FLAG pulldown was repeated until DDK was depleted from the extract. Fractions containing DDK were pooled, and the volume reduced to 0.5 ml using an Amicon spin concentrator (Millipore). The pooled, concentrated eluate was fractionated on a 24 ml Superdex 200 Increase 10/300 GL (GE Healthcare) gel filtration column in buffer A / 300 mM NaCl / 1 mM DTT. Fractions were analyzed by SDS-PAGE and peak fractions containing DDK were pooled and concentrated using Amicon spin concentrator before dialysis against buffer A / 100 mM KOAc / 2 mM β-mercaptoethanol. The concentration of the purified DDK was determined by SDS-PAGE and Coomassie stain using BSA standards. Purified DDK was stored in aliquots a −80°C.

#### Nhp6

Nhp6 was expressed as a N-terminal 6x His-tag fusion protein in *E*.*coli* BL21-CodonPlus (DE3)-RIL cells (Agilent). A colony of cells freshly transformed with plasmid p1035 was grown in 3 L of LB supplemented with 50 μg/mL ampicillin and 34 μg/mL chloramphenicol at 37°C. At OD_600_ ∼ 0.6, 1 mM IPTG was added and the temperature reduced to 4°C. After 1 hour, the temperature was raised to 20°C and the cells were incubated for an additional 16 hours. Cells were collected by centrifugation, rinsed twice with dH_2_O, once with buffer B (50 mM Tris-HCl pH 7.5 / 1 mM EDTA / 10 % glycerol / 10 mM benzamidine / 150 mM NaCl), and resuspended in buffer B supplemented with 1x protease inhibitor cocktail (Pierce) and 1 mM DTT. Cells were lysed by addition of 10 mg lysozyme (Thermo Scientific) and incubation for 30 mins at 4°C followed by sonication. The clear, soluble phase was isolated after centrifugation of the whole-cell lysate in a T-647.5 rotor (Thermo Fisher) at 40,000 rpm for 30 minutes at 4°C. Nhp6 was pulled down with 0.5 ml packed Ni-NTA agarose beads (Qiagen) for 3 hours at 4°C with gentle agitation. Beads were collected and washed with 20 volumes of Buffer B / 1 mM DTT. Protein was eluted with 5 volumes of buffer B / 1 mM DTT / 100 mM imidazole, and eluates were analyzed by SDS-PAGE. Eluate fractions containing Nhp6 were pooled and the volume reduced to 0.5 ml using an Amicon spin concentrator (Millipore). The pooled concentrate was fractionated by gel filtration chromatography using a Superdex 200 10/300 GL (GE Healthcare) column in buffer B / 1 mM DTT. Elution fractions were analyzed by SDS-PAGE and Nhp6 peak fractions were pooled and concentrated using a spin concentrator. Purified Nhp6 was aliquoted, snap-frozen, and stored at −80°C.

#### FACT

FACT complex was purified after overexpression in yeast cells harboring codon-optimized copies of *SPT16* and N-terminally CBP-tagged *POB3* under control of the GAL1,10 promoter (strain YDR 125). Cells were grown in 12 L YPLG at 30°C up to a density of 2 × 10^7^ cells / mL. Protein expression was induced with 2 % galactose for 4 hours. Cells were collected by centrifugation, washed with 25 mM HEPES-KOH pH 7.6 / 1 M sorbitol, and resuspended in 0.5 volumes of Buffer C (25 mM Tris-HCl pH 7.5 / 0.02 % NP-40 substitute / 10 % glycerol) / 100 mM NaCl / 1 mM DTT / 1x protease inhibitor cocktail (Pierce). The cell suspension was pipetted dropwise into liquid nitrogen to generate frozen popcorn and stored at −80°C. Cells were lysed by crushing in a Spex freezer mill with 10 cycles of 2 minute run and 1 minute cooldown at 15 CPS. Thawed whole cell lysate was supplemented with 1 volume of Buffer C / 100 mM NaCl / 1 mM DTT / 1x protease inhibitor cocktail and the final concentration of NaCl adjusted to 300 mM using a 5 M NaCl stock solution. After 20 minutes of gentle agitation at 4°C, the cell lysate was centrifuged in a T-647.5 rotor (Thermo Fisher) at 40,000 rpm for 1 hour at 4°C. The clear soluble phase was recovered and supplemented with 2mM CaCl_2_. FACT was pulled down from the extract with 0.5 ml packed calmodulin affinity resin (Agilent) for 4 hours at 4°C with gentle rocking. Resin was collected by centrifugation and washed with 10 volumes of Buffer C / 300 mM NaCl / 2 mM CaCl_2_ / 1mM DTT. Protein was eluted from the resin with 7 volumes of Buffer C / 300 mM NaCl / 1 mM EDTA / 2 mM EGTA / 1 mM DTT, and eluates were analyzed by SDS-PAGE and Coomassie stain. The calmodulin pulldown was repeated until FACT was depleted from the extract. Fractions containing FACT were pooled and incubated for 16 hours at 4°C with 400μg TEV protease to remove the CBP tag. The digest was diluted 3-fold with buffer D (25 mM Tris-HCl pH 7.5 / 1 mM EDTA / 10 % glycerol) / 1mM DTT to reduce the final NaCl concentration to 100 mM, and fractionated on a MonoQ 5/50 GL (GE Healthcare) column in buffer D / 1mM DTT using a gradient of 0.1 – 1 M NaCl over 20 column volumes. Fractions containing FACT were pooled and concentrated to a volume of 0.5 ml using an Amicon spin concentrator (Millipore). The concentrate was gel-filtered on a 24 ml Superdex 200 10/300 GL (GE Healthcare) column equilibrated in 25 mM HEPES-KOH pH 7.5 / 1 mM EDTA / 10 % glycerol / 300 mM KOAc / 1 mM DTT. Elution fractions were analyzed by SDS-PAGE and Coomassie stain. FACT-containing peak fractions were pooled, spin-concentrated, aliquoted, snap-frozen and stored at - 80°C.

#### Cdt1·MCM^2Δ127^

Purified Cdt1·Mcm2-7^2-TEV^ complex was supplemented with 22.5-fold molar excess of TEV protease and incubated for 1 hour at 30°C. The reaction was fractionated on a Superdex 200 Increase 10/300 GL (GE Healthcare) gel filtration column in 45 mM HEPES-KOH pH 7.6 / 5 mM Mg(OAc)_2_ / 0.02 % NP-40 substitute / 10 % glycerol / 100 mM KOAc / 1 mM ATP / 1 mM DTT, and Cdt1·MCM^2Δ127^-containing peak fractions pooled, aliquoted, snap-frozen in liquid nitrogen and stored at −80°C.

### DNA templates

#### DNA beads

MCM loading, MCM phosphorylation, and MCM-DDK binding assays were performed on a linear 3 kb ARS305-containing DNA covalently linked on one end to HpaII methyltransferase and immobilized on paramagnetic beads via a 5’ photocleavable biotin on the other end. The template was PCR-amplified from p470 using oligo DR772, which contains a photocleavable 5’ biotin moiety, and oligo DR2417, which contains a M.HpaII-binding sequence modified with 5-fluoro-2′-deoxycytidine (BioSynthesis). The purified PCR product was coupled to Dynabeads M280 streptavidin magnetic beads (Invitrogen). M.HpaII (NEB) was conjugated to bead-bound DNA in 50 mM Tris-HCl pH 7.5, 10 mM EDTA, 100 μM SAM at a ratio of 4 units M.HpaII per 90 fmol of DNA for 16 hours at 37°C with agitation. M.HpaII-conjugated bead-bound DNA was washed and stored in 10 mM HEPES-KOH pH 7.6 / 50 mM KOAc / 1 mM DTT at 4°C.

#### Plasmids

The plasmid unwinding assay was performed on circular 3 kb ARS1-containing p79, while the *in vitro* DNA replication assays were performed on ARS1-containing p1017 (4.8 kbp) or ARS305-containing p470 (10 kbp) DNA. Plasmid DNAs were initially isolated using a maxiprep kit (Qiagen). To remove nicked plasmid species, purified plasmid DNA was fractionated on a 10 – 40 % sucrose gradient in 20 mM Tris-HCl pH 7.5 / 1mM EDTA / 1M NaCl using an AH-629 swinging bucket rotor (Thermo Scientific) at 27,000 rpm for 20 hours at 20°C. 0.5 ml fractions were collected and analyzed by agarose gel-electrophoresis in the absence of ethidium bromide. The gel was stained post-run with ethidium bromide. Supercoiled DNA-containing fractions were pooled, dialyzed against 10 mM Tris pH 7.5 / 2mM EDTA, concentrated using an Amicon spin concentrator (Millipore) to 1 to 2 mg/ml, and stored in aliquots at -20°C.

To generate chromatinized templates for DNA replication assays, 1.5 μg Nap1, 0.5 μg Histone octamer, 0.2 μg Isw1a, and 0.8 pmol ORC were mixed and incubated in 10 mM HEPES-KOH pH 7.5 / 50 mM KCl / 5 mM MgCl_2_ / 0.5 mM EGTA / 10 % glycerol / 0.1mg/mL BSA for 30 minutes at 4°C. 0.5 μg of purified supercoiled plasmid DNA was subsequently added along with 3 mM ATP, 20mM creatine phosphate, and 50 ng/μL creatine kinase in a total volume of 10 μL for 1 hour at 30°C. The chromatin template was immediately used for *in vitro* replication.

### MCM loading assay

MCM loading reactions were carried out with agitation for 30 minutes at 30°C in a 40 μL reaction volume in 25 mM HEPES-KOH pH 7.6 / 10 mM Mg(OAc)_2_ / 0.02 % NP-40 substitute / 5 % glycerol / 100 mM KOAc / 1 mM DTT / 5 mM ATP or ATPγS, using 88 nM ORC, 86 nM Cdc6, 420 nM Cdt1·Mcm2-7 (wildtype or mutant as indicated), and 0.3 μg of bead-bound DNA. After the reaction, beads were magnetically separated from the supernatant, and washed once with Wash Buffer (45 mM HEPES-KOH pH 7.6 / 5 mM Mg(OAc)_2_ / 1 mM EDTA / 1 mM EGTA / 0.02 % NP-40 substitute / 10 % glycerol) / 300 mM KOAc, once with Wash Buffer / 500 mM NaCl, and once with Binding Buffer (25 mM HEPES-KOH pH 7.6 / 10 mM Mg(OAc)_2_ / 0.02 % NP-40 substitute / 5% glycerol / 100 mM KOAc). Beads were resuspended in Binding Buffer / 1mM DTT and supplemented with either 9.45 μM TEV protease (a 22.5-fold molar excess over Cdt1·Mcm2-7) or an equal volume of TEV protease storage buffer (50 mM Tris-HCl pH 7.5 / 1 mM EDTA / 10 % glycerol / 100 mM NaCl / 1 mM DTT) as a mock control in a total volume of 40 μL. The reactions were incubated for 1 hour at 30°C with agitation. Beads were magnetically separated from the supernatant and washed once with Wash Buffer / 300 mM KOAc, once with Wash Buffer / 500 mM NaCl, and once with Binding Buffer. Beads were resuspended in 20 μL Wash Buffer / 50 mM KOAc / 1 mM DTT, and the DNA eluted from the beads by exposure to UV_312 nm_ for 10 minutes using a hand-held UV lamp. The supernatant, containing the DNA and DNA-bound proteins, was analyzed by SDS-PAGE followed by silver staining.

### MCM phosphorylation assay

MCM loading was carried out as described above. Following the TEV protease or mock cleavage and wash steps, beads were resuspended in Binding Buffer / 5mM ATP / 1mM DTT and supplemented with purified DDK at 150 nM or indicated concentrations in a total volume of 40 μL. The reaction was incubated for 20 minutes at 30°C with agitation. Beads were magnetically separated from the supernatant, and washed once with Wash Buffer / 300 mM KOAc, once with Wash Buffer / 500 mM NaCl, and once with Binding Buffer. Beads were resuspended in 20 μL Wash Buffer / 50 mM KOAc / 1 mM DTT, and the DNA was eluted from the beads by exposure to UV_312 nm_ for 10 minutes. The supernatant, containing the DNA and DNA-bound proteins, was analyzed by SDS-PAGE and silver staining.

### DDK-MCM DH binding assay

MCM loading was carried out as described above. Following the TEV protease or mock cleavage and wash steps, beads were resuspended in Binding Buffer / 5 mM ATPγS / 1mM DTT and supplemented with purified DDK at 150 nM or indicated DDK concentrations in a total volume of 40 μL. For Figures 4E, 6A and 6B, ATPγS was substituted with the indicated ATP analog. Binding reactions were incubated for 20 minutes at 30°C with agitation. For Figures 6A & 6B, the indicated Rad53 variant was either pre-mixed with DDK or added sequentially. To pre-mix, 250 nM Rad53 was incubated with 150 nM DDK in 10 mM Mg(OAc)_2_ and 5 mM ATP or AMP-PNP for 20 minutes at 30°C before adding to the resuspended beads. For sequential addition, the binding reaction was carried out as above for 10 minutes instead of 20 minutes, at which point 250 nM Rad53 was added and the reaction carried out for another 10 minutes. Beads were magnetically separated from the supernatant and washed once with Wash Buffer /100 mM KOAc. For Figure 4C, another round of TEV or mock cleavage was performed in Binding Buffer / 5mM ATPγS / 1mM DTT for 1 hour at 30°C followed by a wash with Wash Buffer / 100mM KOAc. For Figure 4D, beads were washed in Wash Buffer with the indicated salt concentrations. Beads were resuspended in 20 μL Wash Buffer / 50 mM KOAc /1 mM DTT, and the DNA was eluted from the beads by exposure to UV_312 nm_ for 10 minutes. The supernatant, containing the DNA and DNA-bound proteins, was analyzed by SDS-PAGE and silver staining.

### Plasmid unwinding assay

0.5 μg of supercoiled p79 plasmid DNA was first relaxed with 100 nM purified Top1 in 25 mM HEPES-KOH pH 7.6 / 10mM Mg(OAc)_2_ / 0.02 % NP-40 substitute / 5% glycerol / 100 mM KOAc / 1 mM DTT in a reaction volume of 10 μL for 1 hour at 30°C. MCM loading was subsequently carried out in 25mM HEPES-KOH pH 7.6 / 10 mM Mg(OAc)_2_ / 0.02 % NP-40 substitute / 5 % glycerol / 100 mM KOAc / 1mM DTT / 3.5 mM ATP using 65 nM ORC, 104 nM Cdc6, 130 nM Cdt1·Mcm2-7, 20 mM creatine phosphate, and 50 ng/μL creatine kinase in a reaction volume of 20 μL for 30 minutes at 30°C. Where indicated, 2.9 μM TEV protease was added (a 22.5- fold molar excess over Cdt1·Mcm2-7) for 1 hour at 30°C. 60 nM DDK was then added for 20 minutes at 30°C. For CMG assembly and activation the reaction was supplemented with 34 nM CDK, 0.5 mg BSA, 40 nM Sld3·7, 40 nM Cdc45, 50 nM GINS, 34 nM Pol ε, 54 nM Dpb11, 40 nM Sld2, 100 nM RPA, 20 nM Top1 and 14 nM Mcm10 in a total volume of 50 μL with the salt concentration adjusted to 185 mM KOAc for 30 minutes at 30°C. The reaction was quenched with 17 mM EDTA, 0.2 % SDS, and 0.8 U Proteinase K (NEB) for 30 minutes at 37°C. DNA was purified by phenol:chloroform extraction and centrifugation through a G-25 spin column (GE Healthcare). Samples were analyzed by agarose gel-electrophoresis and post-run ethidium-bromide stain.

### *In vitro* DNA replication assay

MCM loading was performed for 30 minutes at 30°C in a reaction volume of 20 μL including 40 nM ORC, 64 nM Cdc6, 80 nM Cdt1·Mcm2-7, 0.5 μg plasmid DNA and a reaction buffer containing 25 mM HEPES-KOH pH 7.6 / 10 mM Mg(OAc)_2_ / 0.02 % NP-40 substitute / 5 % glycerol / 100 mM KOAc / 1 mM DTT / 3.5 mM ATP / 20mM creatine phosphate / 50 ng/μL creatine kinase. Where indicated, 1.8 μM TEV protease or an equivalent volume of TEV storage buffer was added for 1 hour at 30°C. 60 – 140 nM DDK was then added for 20 minutes at 30°C. Origin firing was then induced by supplementing the reaction with 0.5 mg BSA, 40 nM Sld3·7, 40 nM Cdc45, 35 nM CDK, 50 nM GINS, 34 nM Pol ε, 30 nM Dpb11, 40 nM Sld2, 135 nM RPA, 40 nM Pol α, 35 nM Ctf4, 40 nM RFC, 35 nM PCNA, 4 nM Pol δ, 14 nM Csm3-Tof1, 14 nM Mrc1, 20 nM Top1, 15 nM Top2, 7 nM Mcm10, 122 μM each NTP, 40 μM each dNTP, and trace amount of α-^32^P-dATP or α-^32^P-dCTP in a total volume of 50 μL with the salt concentration adjusted to 185 mM KOAc for 30 minutes at 30°C. Reactions on chromatin templates were additionally supplemented with 2 μM Nhp6 and 200 nM FACT. The reactions were quenched with 17 mM EDTA, 0.2 % SDS, and 0.8U Proteinase K (NEB) for 30 minutes at 37°C. DNA was isolated by phenol:chloroform extraction and centrifugation through a G-25 spin column (GE Healthcare). Samples were fractionated on 0.8 % denaturing agarose gel in 30 mM NaOH / 2 mM EDTA, dried, and exposed to a phosphorimager screen. Images were scanned ion a Typhoon FLA-9500 and analyzed with ImageJ.

## Acknowledgments

This work was supported by NIGMS grant R01-GM127428.

---

**Supplementary Figure 1.**
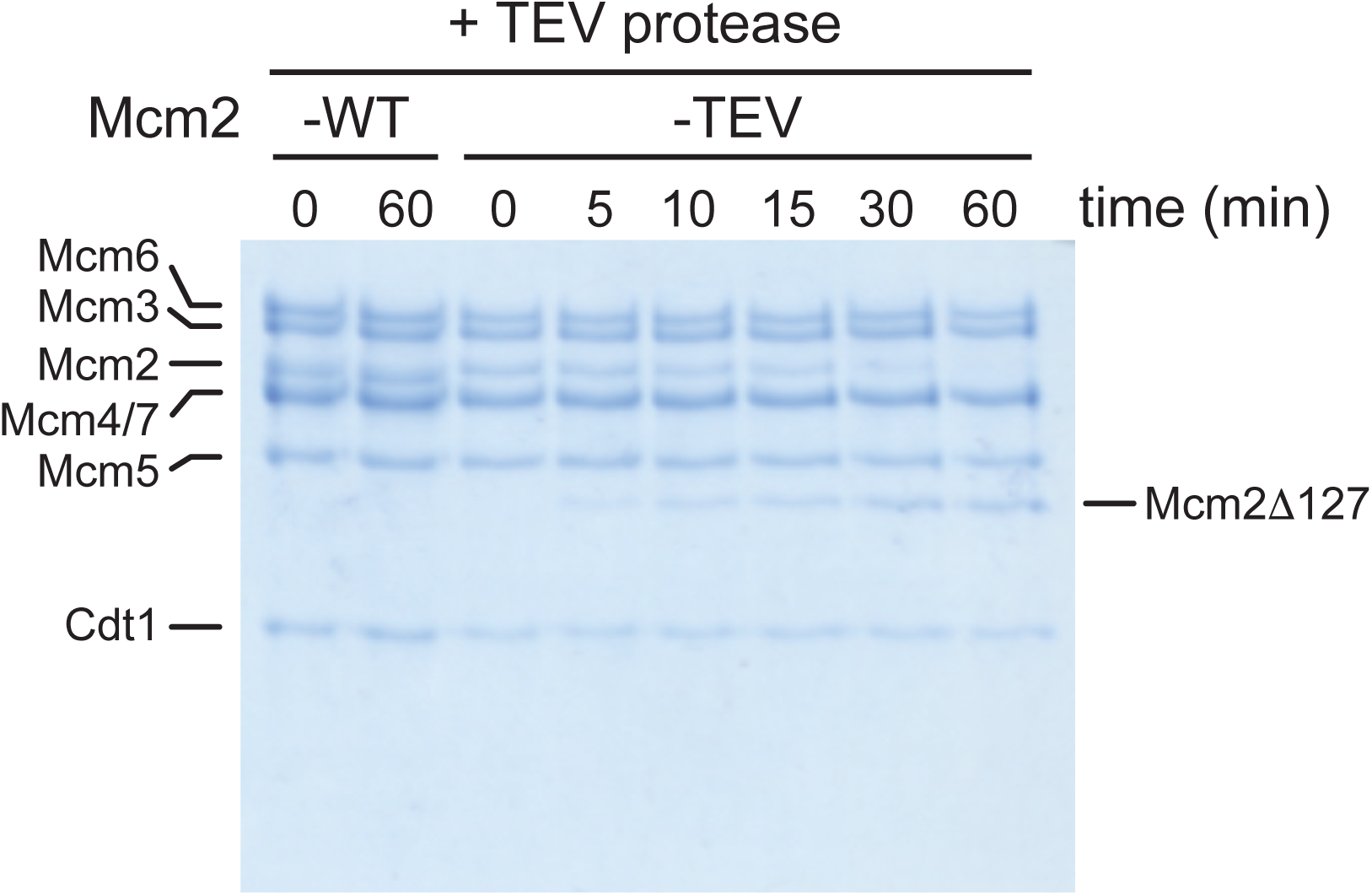
Time course analysis of Cdt1·MCM^2-TEV^ and Cdt1·MCM^2-WT^ cleavage by TEV protease. Fractions of the reactions were analyzed by SDS-PAGE and Coomassie stain.

**Supplementary Figure 2.**
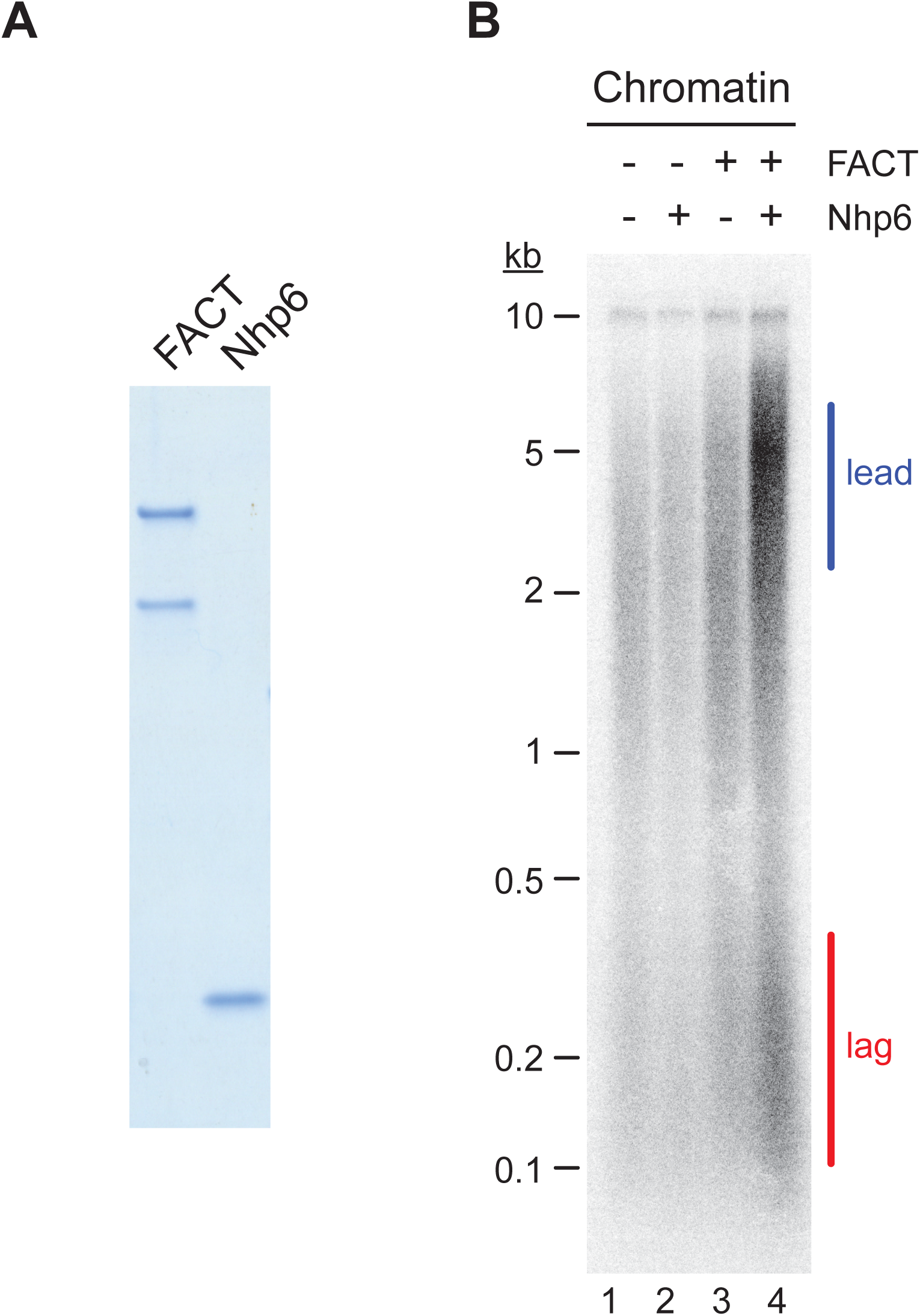
FACT / Nhp6-dependent chromatin replication. (**A**) Purified FACT and Nhp6. Samples were analyzed by SDS-PAGE and Coomassie stain. (**B**) *In vitro* DNA replication reaction was performed on chromatinized p470 (10 kbp) in the absence or presence of FACT and Nhp6 as indicated. Reaction products were analyzed by denaturing agarose gel-electrophoresis and autoradiography.

**Supplementary Figure 3.**
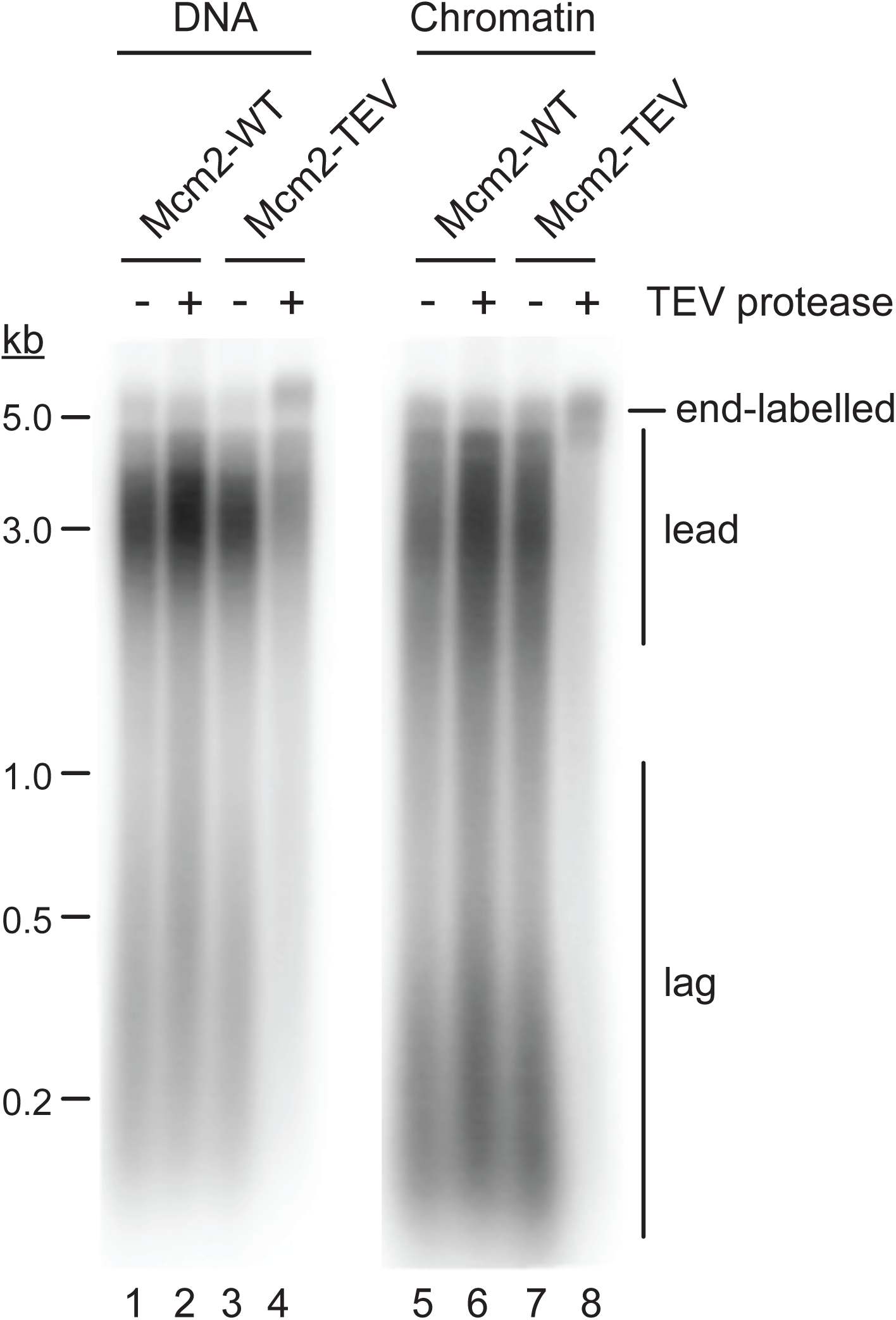
Attenuation of DNA synthesis in the presence of Mcm2-TEV is dependent on TEV protease cleavage. Standard DNA replication reactions using naked or chromatinized p1017 (4.8 kbp) as template were performed with Cdt1·MCM^2-TEV^ or Cdt1·MCM^2-WT^ as indicated. TEV protease or mock buffer was added to the reaction following Mcm2-7 loading and preceding origin activation as indicated. Reaction products were analyzed by denaturing agarose gel-electrophoresis and autoradiography.

**Supplementary Figure 4.**
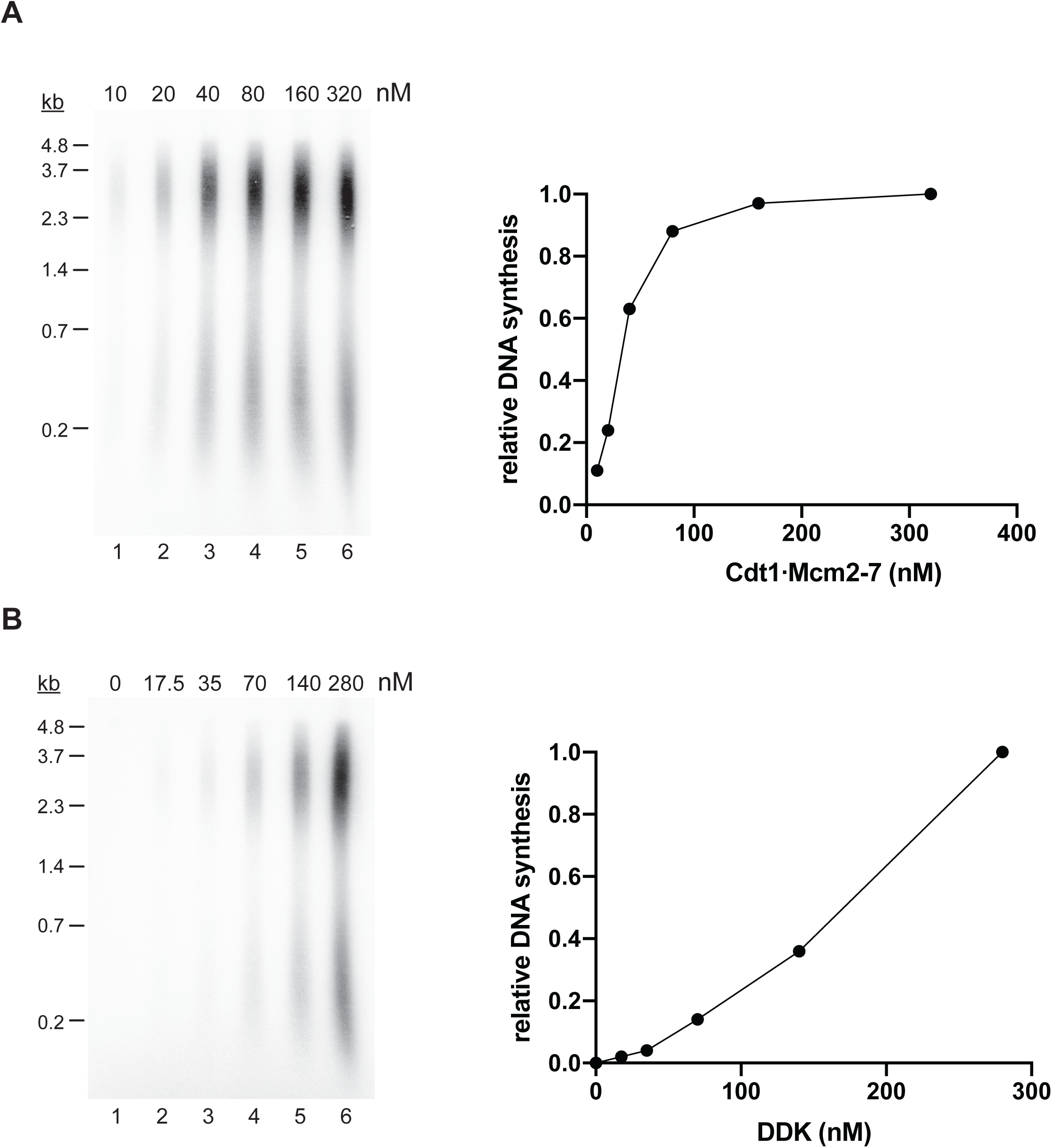
(**A**) Cdt1·Mcm2-7 titration experiment using standard DNA replication conditions. Template: p1017 (4.8 kbp). Left: Reaction products were analyzed by denaturing agarose gel-electrophoresis and autoradiography. Right: Plot of total normalized DNA synthesis. (**B**) DDK titration experiment using standard DNA replication conditions, but Cdt1·MCM^2-Δ127^ in place of Cdt1·MCM^2-WT^. Template: p1017 (4.8 kbp). Left: Reaction products were analyzed by denaturing agarose gel-electrophoresis and autoradiography. Right: Plot of total normalized DNA synthesis.

